# How do symbiotic associations in lecideoid lichens respond to different environmental conditions along the Transantarctic Mountains, Ross Sea region, Antarctica?

**DOI:** 10.1101/2021.05.26.445136

**Authors:** Monika Wagner, Georg Brunauer, Arne C. Bathke, S. Craig Cary, Roman Fuchs, Leopoldo G. Sancho, Roman Türk, Ulrike Ruprecht

## Abstract

Lecideoid lichens as dominant vegetation-forming organisms in the climatically harsh areas of the southern part of continental Antarctica show clear preferences in relation to environmental conditions (i.e. macroclimate). 306 lichen samples were included in the study, collected along the Ross Sea coast (78°S–85.5°S) at six climatically different sites. The species compositions as well as the associations of their two dominant symbiotic partners (myco- and photobiont) were set in context with environmental conditions along the latitudinal gradient. Diversity values were nonlinear with respect to latitude, with the highest alpha diversity in the milder areas of the McMurdo Dry Valleys (78°S) and the most southern areas (Durham Point, 85.5°S; Garden Spur, 84.5°S), and lowest in the especially arid and cold Darwin Area (~79.8°S). Furthermore, the specificity of mycobiont species towards their photobionts decreased under more severe climate conditions. The generalist lichen species *Lecanora fuscobrunnea* and *Lecidea cancriformis* were present in almost all habitats, but were dominant in climatically extreme areas. *Carbonea vorticosa, Lecidella greenii* and *Rhizoplaca macleanii* were confined to milder areas.

In summary, the macroclimate is considered to be the main driver of species distribution, making certain species useful as bioindicators of climate conditions and, consequently, for detecting climate change.

## Introduction

Polar deserts of the southernmost areas in continental Antarctica are characterized by exceptionally hostile climatic conditions, such as particularly low temperatures and high aridity (Adams et al. 2006; Cary et al. 2010; Magalhaes et al. 2012).Terrestrial life is restricted to ice-free areas, which, apart from a few nunataks, are mainly located along the Transantarctic Mountains forming the west coast of the Ross Sea and Ross Ice Shelf (Monaghan et al. 2005). Because of these special conditions, terrestrial life is rare and can only be found in small areas protected from extreme environmental influences, such as abrasion from windblown particles or high solar radiation, the so-called microhabitats (Hertel 1998; Ruprecht et al. 2012b). They are characterized by sheltered areas in rock crevices or small cavities shielded from the wind and sun that allow life on a small scale in an otherwise hostile environment. The rock surface is often highly weathered which results in a higher water retention capacity, providing the most needed life source for the organisms to survive (Colesie et al. 2014; Green 2009). The only moisture available to rock-dwelling organisms is provided by clouds, fog, dew, sparse precipitation and melting snow (Head and Marchant 2014; Wagner et al. 2020). Additionally, the aspect of the slopes, ridges and depressions as well as the wind regime has an important impact by creating different surface temperatures in small areas (McKendry and Lewthwaite 1990; Yung et al. 2014). However, microhabitats are influenced by both macroclimate and geography, and their life-supporting properties therefore vary along environmental gradients reflected in changing diversity levels and biogeography of Antarctic terrestrial biota (Baird et al. 2019; Lagostina et al. 2021; Magalhaes et al. 2012; Peat et al. 2007; Fig.1; Ruprecht et al. 2012a; Fig.1).

**Figure 1.**
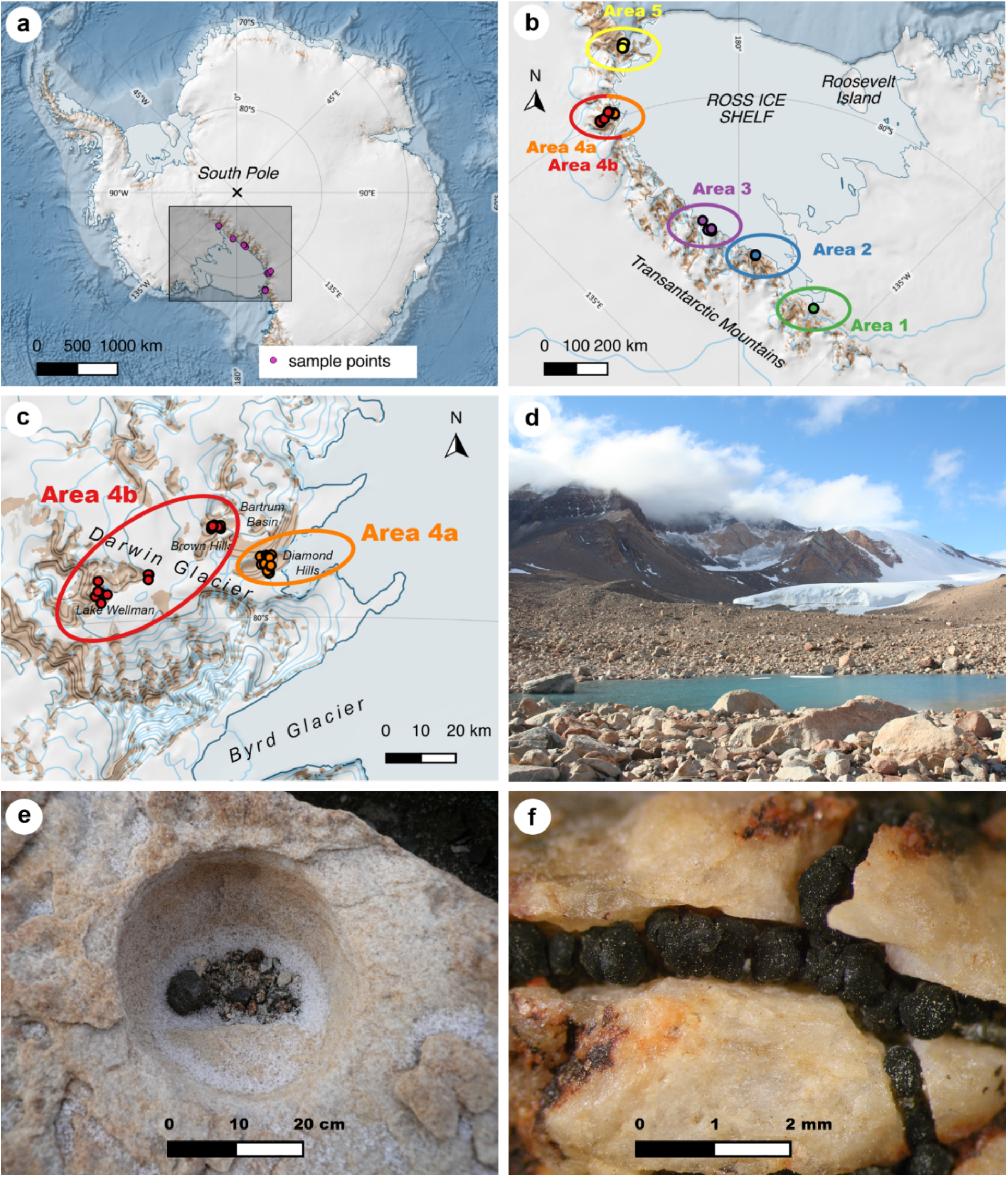
Location of the sample points and lichen habitats. (a) Antarctic continent, investigated area marked with rectangle, (b) location of the six different areas defined in the study, (c) differentiation of area 4 in subareas 4a and 4b, (d) Batrum Basin, (e) microhabitat with crustose lichens at Lake Wellman, (f) chasmolithic growth of *Lecidea cancriformis*. Maps of (a), (b) and (c) are based on the dataset Quantarctica (Matsuoka et al. 2018).

The terrestrial vegetation along the Ross Sea coast (extending from 72°S, Cape Hallett, to 85.5°S, Queen Maud Mountains) is entirely composed of cryptogrammic organisms and dominated by lichens and mosses (Colesie et al. 2014; Ochyra et al. 2008; Peat et al. 2007). Remarkably, the biodiversity of these organisms does not decrease evenly along the latitudinal gradient as one might expect. In fact, the lowest species diversity was recorded at about 79°S at Diamond Hill (Darwin Area), which has by far the harshest climate conditions (lowest humidity; Colesie et al. 2014). However, nonlinear climatic conditions along gradients caused by additional factors, e.g. special wind systems, can be detected effectively with biological systems that act as bioindicators (Dal Grande et al. 2017; Sancho et al. 2019; Singh et al. 2017; Wagner et al. 2020). Additionally, they not only enable the survey of the current state but can also reliably indicate changes in environmental conditions. Due to the structure and diversity of communities, the abundance and distribution of species as well as processes varying along environmental gradients are therefore powerful long-term and large-scale study systems to estimate the consequences of climate change on ecosystems (Sundqvist et al. 2013).

The most abundant vegetation-forming organisms in these areas are lichens, in most of the cases with a crustose thallus fused to the rocky surface or deeply embedded in crevices (Colesie et al. 2014; De los Rios et al. 2004; Hertel 2007; Kappen and Valladares 2007; Ruprecht et al. 2012b). The poikilohydric lifestyle of lichens enables them to survive the harsh climate conditions and the long periods without water and/or light in a dormant state (Schroeter et al. 2011). The symbiotic lifeform of lichens consists of two dominant symbiotic partners: the mycobiont (fungus) and the photosynthetic partner (green algae and/or cyanobacteria: photobiont) and additional associated fungal, algal and bacterial communities forming the holobiome lichen thallus (Aschenbrenner et al. 2016; Grube et al. 2015; Lawrey and Diederich 2003; Ruprecht et al. 2014; Spribille et al. 2016). Therefore, lichens constitute an excellent model for analyzing multi-species associations in one unit to reveal phylogenetic and ecological responses for symbiotic associations.

Many analyses focused on myco-/photobiont associations have demonstrated that they react sensitively to even small environmental gradients (Dal Grande et al. 2018; Wagner et al. 2020). These results allow the conclusion that mycobionts which are less specialized to specific locations and are able to use a broader range of photobionts, such as the widespread species *Lecidea cancriformis* in continental Antarctica (Ruprecht et al. 2012a; Wagner et al. 2020), are less vulnerable to climate changes. Low photobiont specificity may improve the performance of the lichen symbiosis, e.g. by increasing the adaptive potential to new environmental conditions, and widening the geographical range via ecological niche shifts (Leavitt et al. 2015; Rolshausen et al. 2018; Vančurová et al. 2018). On the other hand, high levels of photobiont specificity are expected under conditions where ecological factors, especially (macro-) climate and/or substrate (e.g. calcareous or siliceous rock), exert a strong selective influence on lichen performance (Peksa and Skaloud 2011; Vančurová et al. 2018; Werth and Sork 2010). Additionally, genetic identity can play a significant role in shaping myco-/photobiont associations along gradients (Dal Grande et al. 2017) or may also lead to turnover zones, suggesting that photobionts are replaced by others as environmental conditions change (Rolshausen et al. 2020). An influence on the selection of *Trebouxia* species due to temperature combined with water availability was suggested in several studies as a key factor of photobiont selection of lichens in Antarctica (Green et al. 2011a; Wagner et al. 2020). Due to the sensitive response of lichen communities to climatic change with modified species compositions and reduced diversity (Ellis 2019; Mayer et al. 2013; Sancho et al. 2019; Sancho et al. 2017) lichen growth, abundance and diversity are expected to be negatively affected by climatic changes (Sancho et al. 2017). Consequently, lichens represent excellent bioindicators because of their sensitive responses to environmental changes (Alatalo et al. 2015; Allen and Lendemer 2016; Bassler et al. 2016; Sancho et al. 2019), and especially abundant and cosmopolitan species serve as a valuable model system to record diversity and composition along climatic gradients worldwide.

The current study focuses on the association patterns of the two main symbionts (myco- and photobiont) of the lecideoid lichen group (Ruprecht et al. 2020) that is dominant along the investigated part of the latitudinal gradient (78–85°S) at the Ross Sea coast. The following objectives were addressed: (1) to assess the biodiversity and genetic identity of the symbiotic partners of lecideoid lichens using phylogenic methods; (2) to investigate how the variability of myco-/photobiont associations is related to environmental variables (elevation, temperature, precipitation) using diversity and specificity indices as well as network statistics, and (3) to identify certain myco-/photobiont associations that are representative for climatic conditions and therefore may qualify as bioindicators.

## Materials & Methods

### Study area and investigated lichen specimens

The sample sites were divided in five different main areas (Fig. 1b). Area 4 (Darwin Area) was then subdivided in subareas 4a and 4b, considering the wide range of climate conditions within this region (Figure 1c). Site descriptions of the six regions are given in Table 1, geographical descriptions can be found at the Supplementary Table S1.

**Table 1.** Site descriptions of the six regions defined in the present study, including range of the coordinates of the sampling sites and areas, the number of sampling sites and the BIOCLIM variables BIO10 (mean temp. of the warmest quarter) and BIO12 (annual precipitation) per area.

|  | Sampling area | Range of coordinates of sampling sites | Number of sampling sites | Elevation mean (m. a. s. l.) | BIO10: mean temperature, warmest quarter (°C) | BIO12: precipitation, annual mean (mm) |
| --- | --- | --- | --- | --- | --- | --- |
| **Area 1** | Scott Glacier/<br>Durham Point | S 85.54°<br>W 151.15° | 1 | 370.00 | -7.30 | 190.00 |
| **Area 2** | Massam Glacier/<br>Garden Spur | S 84.54°–84.56°<br>W 174.91°–175.01° | 2 | 182.92 | -6.96 | 113.00 |
| **Area 3** | Mt. Kyffin, The<br>Gateway, Mt.<br>Harcourt | S 83.49°–83.83°<br>E 170.79°–172.76° | 6 | 774.72 | -8.20 | 104.37 |
| **Area 4a** | Darwin Area:<br>Diamond Hills,<br>Brown Hills | S 79.84°–79.88°<br>E 159.22°–159.39° | 30 | 484.19 | -8.23 | 91.79 |
| **Area 4b** | Darwin Area:<br>Bartrum Basin,<br>Smith Valley, Lake<br>Wellman | S 79.75°–79.95°<br>E 156.70°–158.67° | 31 | 726.39 | -10.18 | 69.13 |
| **Area 5** | McMurdo Dry<br>Valleys | S 78.02°–78.17°<br>E 163.62°–164.10° | 102 | 589.42 | -6.24 | 145.04 |

Altogether 306 lecideoid lichen specimens were collected on siliceous substrate along a latitudinal gradient (78–85.5°S) of the southwest Ross Sea coast (Antarctica, Fig.1a-c). 147 samples of the genera *Carbonea, Lecanora, Lecidella* and *Lecidea* were collected at 70 different localities from the following sampling areas: Area 1, Scott Glacier/Durham Point; Area 2, Massam Glacier/Garden Spur; Area 3, Mt. Kyffin, Mt. Harcourt, The Gateway; Area 4a, Darwin Area (Diamond Hills, Brown Hills); Area 4b, Darwin Area (Bartrum Basin, Smith Valley and Lake Wellman) (Fig. 1a-c, Supplementary Table S1). To get a better coverage of the latitudinal gradient (Fig. 1a), additionally, 159 lichen samples (collected at 102 different localities) from the Area 5, McMurdo Dry Valleys (MDV), were obtained from the studies of Wagner et al. (2020) and Perez-Ortega et al. (2012), including the solely lecideoid lichen species of the genera *Carbonea, Lecanora, Lecidella, Lecidea* and *Rhizoplaca*. The entire lists of samples can be found at the Supplementary Tables S2–4.

All voucher specimens are stored in the herbarium of the University of Salzburg (SZU) except for samples collected by Leopoldo G. Sancho which are deposited in the MAF herbarium of the Botany Unit, Fac. Farmacia, in Madrid.

### DNA-amplification, primer-design and sequencing

Total DNA was extracted from individual thalli by using the DNeasy Plant Mini Kit (Qiagen) following the manufacturer’s instructions. For all samples, the internal transcribed spacer (ITS) region of the mycobionts’ and photobionts’ nuclear ribosomal DNA (nrITS) were sequenced and amplified. Also, additional markers were amplified: for the mycobionts the mitochondrial small subunit (mtSSU) and the low-copy protein coding marker *RPB1*; for the photobionts, the chloroplast-encoded intergenic spacer (psbJ-L) and part of the cytochrome oxidase subunit 2 gene (COX2). This was done using specific primers and PCR-protocols in our project framework (Ruprecht et al. 2020).

The nrITS of the mycobiont was amplified using the primers ITS1F (Gardes and Bruns 1993), ITS4(White et al. 1990), ITS1L (Ruprecht et al. 2020) and ITS4L (Ruprecht et al. 2020). The nrITS of the photobiont was amplified using the primers 18S-ITS uni-for (Ruprecht et al. 2012a), ITS4T (Kroken and Taylor 2000), ITS1T (Kroken and Taylor 2000) and ITS4bT_mod (5’-CCAAAAGGCGTCCTGCA-3’; modified, based on Ruprecht et al. (2014)). For the marker mtSSU, the primer mtSSU rev2 (Ruprecht et al. 2010) and the newly designed primers mtSSU for2 mod1 (5’-AACGGCTGAACCAGCAACTTG-3’) and mtSSU rev1 (5’-AGGYCATGATGACTTGTCTT-3’) were used. For *RPB1*, gRPB1-A for (Matheny et al. 2002), fRPB1-C rev (Matheny et al. 2002) and RPB1_for_Lec (Ruprecht et al. 2020) were chosen. For the marker COX2, COXIIf2 and COXIIr (Lindgren et al. 2014) and COXII_sense (Ruprecht et al. 2020) were used, for psbJ-L, newly designed psbL_for1 (5’-GTTGAATTAAATCGTACTAGT-3’) psbL-sense and psbJ-antisense (Ruprecht et al. 2014) were chosen.

Unpurified PCR-products were sent to Eurofins Genomics/Germany for sequencing.

### Phylogenetic analysis

For both symbionts, the sequences were assembled and edited using Geneious version 8.0.5 (https://www.geneious.com) and aligned with MAFFT v7.017 (Katoh et al. 2002).

Maximum likelihood analyses were calculated with IQ-TREE v1.6.12 (Nguyen et al. 2014), using the model selection algorithm ModelFinder (Kalyaanamoorthy et al. 2017). Branch supports were obtained with the implemented ultrafast bootstrap (UFBoot; Minh et al. 2013). Number of bootstrap alignments: 1000, maximum iteration: 1000, minimum correlation coefficient: 0.99. Additionally, a SH-aLRT branch test (Guindon et al. 2010) was performed. Each branch of the resulting tree was assigned with SH-aLRT as well as UFBoot supports. The branches with SH-aLRT < 80 % and/ or UFboot < 95 % were collapsed by adding the command -minsupnew 80/95 to the script.

In order to be able to use all samples with an incomplete marker set, a multi-marker phylogeny with a reduced number of samples and, in comparison, the complete data set with the marker ITS were calculated and compared, respectively for each symbiont.

For the photobiont, the classification and labeling of the different operational taxonomical units (OTUs) followed the concepts of Muggia et al. (2020) and Ruprecht et al. (2020), using automatic barcode gap discovery (ABGD; Puillandre et al. 2012), based on the marker ITS. The threshold of 97.5 % sequence similarity set by Leavitt et al. (2015) and applied by Ruprecht et al. (2020) was used to ensure clear delimitation of OTUs and sub-OTUs.

### Analysis of spatial distribution

Unless stated otherwise, analysis was conducted in R (R Core Team 2020; version 3.6.3, https://www.r-project.org) using RStudio (RStudio Team 2016; version 1.1.463, https://rstudio.com); figures where produced using the R package ggplot2 (Wickham 2009) and processed using Adobe Photoshop (version 22.2.0., https://www.adobe.com).

Based on data from CHELSA (*Climatologies at high resolution for the earth’s land surface areas*; Karger et al. 2017), the 19 BIOCLIM variables (Nix 1986) were calculated for each sample point using the R functions raster() and extract() of the package raster (Hijmans 2020). These variables are derived variables from the monthly minimum, maximum, mean temperature and mean precipitation values, developed for species distribution modeling and related ecological applications (Karger et al. 2017). For the analyses of this study, BIO10 (mean temperature of the warmest quarter) and BIO12 (annual precipitation) were chosen, as these two variables showed the strongest correlations with the diversity and specificity indices (see below).

For analyzing the spatial distribution of the lichen samples, alpha, beta and gamma diversity values were calculated. The concept was developed in 1960 by Whittaker (Whittaker 1960) who distinguished three aspects or levels of species diversity in natural communities: (1) alpha diversity, the species richness within a particular area, (2) beta diversity, the extent of changes in species diversity between the areas, and (3) gamma diversity, a measure of the overall diversity for the different areas within the whole region. These diversity indices were calculated separately for mycobiont species and photobiont OTUs, using the R functions AlphaDiversity(), BetaDiversity() and GammaDiversity() of the package entropart (Marcon and Hérault 2015), which give reduced-bias diversity values (diversity order: *q = 1* (Shannon diversity); weights: *w_i_ = n_i_/n* with *n_i_*, number of samples in area *i* and *n*, total number of samples). Next, alpha diversity was analyzed for correlations with the following variables: elevation, latitude, BIO10 and BIO12.

To determine whether mycobiont species or photobiont OTU community composition are related to environmental variables (elevation, BIO10 and BIO12), constrained analyses of principal coordinates were conducted, using the R function capscale() of the package vegan (Oksanen et al. 2019; distance: Bray Curtis). Prior to analysis, to standardize species composition data (convert species abundances from absolute to relative values), a Hellinger transformation was performed on the community matrix, using the R function decostand() of the package vegan (Oksanen et al. 2019). The variance explained by constrained ordination was tested by a Monte Carlo permutation test, using the R function anova() of the package vegan (Oksanen et al. 2019).

A Mantel test was performed to test whether the differences in mycobiont species and photobiont OTU community composition between samples are related to physical distance, using the R function mantel() of the package vegan (Oksanen et al. 2019).

### Haplotype analysis

In order to ensure that the entire data set could be processed, all further analyses were carried out using only complete sequences of the marker ITS for all calculations. The number of haplotypes, *h,* of the different mycobiont species and photobiont OTUs was determined using the function haplotype() of the R package pegas (Paradis 2010). Haplotype networks were computed, using the function haploNet() of the R package pegas (Paradis 2010) for mycobiont species and photobiont OTUs with *h* ≥ 2 and at least one haplotype with *n* ≥ 3 (*Carbonea* sp. 2, *Lecanora fuscobrunnea*, *Lecidea cancriformis, Lecidella greenii, Lecidella siplei, Lecidella* sp. nov2 and *Rhizoplaca macleanii,* as well as *Tr*_A02, *Tr*_I01 and *Tr*_S02). The frequencies were clustered in 10% ranges, for example the circles of all haplotypes making up between 20-30% have the same size. Additionally, for the most common mycobiont *L. cancriformis* and the photobiont OTU *Tr*_S02, haplotype networks based on multimarker data sets were calculated, to show that the distribution of haplotypes remains congruent.

### Diversity and specificity indices of mycobiont species and photobiont OTUs

The haplotype as well as the nucleotide diversity was calculated for each identified mycobiont and photobiont species with more than one sample, using the functions hap.div() and nuc.div() of the R package pegas (Paradis 2010), respectively. The haplotype diversity, *Hd,* represents the probability that two randomly chosen haplotypes are different (Nei 1987), the nucleotide diversity, *π*, gives the average number of nucleotide differences per site between two randomly chosen DNA sequences (Nei and Li 1979). Additionally, the ratio of the number of haplotypes *h* divided by the number of samples *N* was calculated.

Furthermore, different metrics for quantifying the phylogenetic species diversity and the specificity of the mycobiont species and photobiont OTUs towards their interaction partners were calculated. Those included the indices *NRI* (Net relatedness index), *PSR* (Phylogenetic species richness) and the Pielou evenness index *J’*. *(Note: to make interpretation similar to the other metrics, for further analyses* 1 – J’ *instead of* J’ *was used.)* An overview of these diversity metrics is given in Supplementary Table S5.

In order to analyze the correlation of these diversity metrics with environment, for every mycobiont species and photobiont OTU with *n* ≥ 10 (*Carbonea* sp. 2, *C. vorticosa, Lecanora fuscobrunnea, Lecidea cancriformis, L. polypycnidophora, Lecidella greenii*, *L. siplei, Rhizoplaca macleanii* and *Trebouxia* OTUs *Tr*_A02, *Tr*_I01, *Tr*_S02, *Tr*_S15, *Tr*_S18) the mean values of the sample locations of the following variables were calculated: elevation, latitude, BIO10 and BIO12.

### Analysis of mycobiont – photobiont associations

To analyze the associations between mycobiont species and photobiont OTUs, bipartite networks were computed, using the R function plotweb() of the package bipartite (Dormann et al. 2008). This was done for each area separately. Additionally, for each bipartite network, the index *H_2_’* was calculated. *H_2_’* is derived from Shannon entropy and characterizes the degree of complementary specialization of partitioning among the two parties of the network. It ranges from 0 for the most generalized to 1 for the most specialized case und was computed using the R functions H2fun() of the package bipartite (Dormann et al. 2008).

Usually, in the context of bipartite networks, also the *d’* value (specialization index) is computed. This value was originally defined for pollination networks and calculates how strongly a species deviates from a random sampling of interacting partners available (Dormann 2011). Thus, in the case of lichens, the *d’* value of a mycobiont species is based on the assumption that for every site of a sampling area, the whole set of photobiont OTUs basically is available. As this is not true for this study, this index was not included.

## Results

### Phylogenetic analysis

For both the mycobiont and photobiont molecular phylogenies from multi-locus sequence data (nrITS, mtSSU and *RPB1* for the mycobiont (140 samples) and nrITS, psbJ-L and COX2 for the photobiont (139 samples) were inferred (Supplementary Fig. S1 and S3). Additionally, phylogenies based solely on the marker nrITS were calculated (Supplementary Fig. S2 and S4), to include samples where the additional markers were not available. Both analyses include only accessions from the study sites (Fig. 1, Table 1). The phylogenies based on the multi-locus data were congruent to the clades of the phylogenies based on the marker nrITS. Thus, in the following, the focus will be only on the latter.

#### Mycobiont

The final data matrix for the phylogeny based on the marker nrITS comprised 306 single sequences with a length of 550 bp. It included sequences of the families *Lecanoraceae* and *Lecideaceae*. The phylogenetic tree was midpoint rooted and shows a total of 19 strongly supported clades on species level, assigned to five genera. The backbone is not supported and therefore the topology will not be discussed. All genera are clearly assigned to their family level and are strongly supported. Only *Lecanora physicella* forms an extra clade as sister to the families *Lecideaceae* and *Lecanoraeae*, which is not the case at the multimarker phylogeny. *L. physciella* has still an uncertain status, because of morphological similarities to both sister families (Ruprecht et al. 2012b). The clade of the genus *Lecidea* revealed seven species (*L. andersonii*, *L. polypycnidophora*, *L.* UCR1, *L.* sp. 5, *L. lapicida*, *L. cancriformis* and *L.* sp. 6), *Lecanora* five species (*L. physciella*, *L.* sp. 2, *L. fuscobrunnea*, *L.* cf. *mons*-*nivis*, *L.* sp. 3), *Carbonea* three species (*C.* sp. URm1, *C. vorticosa*, *C.* sp. 2), and *Lecidella* three species (*L. greenii*, *L. siplei*, *L.* sp. nov2). The samples allocated to the genus *Rhizoplaca* were monospecific (*R. macleanii*). The taxonomical assignment of the obtained sequences were based on the studies of Ruprecht et al. (2020) and Wagner et al. (2020).

#### Photobiont

The final data matrix for the phylogeny based on the marker nrITS comprised 281 single sequences with a length of 584 bp. The phylogenetic tree was midpoint rooted and shows six strongly-supported clades, assigned to seven different OTU levels (Puillandre et al. 2012), using the concept of Muggia et al. (2020) and Ruprecht et al. (2020). The backbone is not supported and therefore the topology will not be discussed. All of the OTUs belong to the genus *Trebouxia* (clades A, I, S), comprising *Tr_*A02, *Tr*_A04a, *Tr*_I01, *Tr*_I17, *Tr_*S02, *Tr_*S15 and *Tr_*S18. Photobiont sequences taken from Perez-Ortega et al. (2012), which were labelled only with numbers, were renamed to assign them to the appropriate OTUs (Ruprecht et al. 2020).

### Analysis of spatial distribution

In general, the most common mycobionts were *Lecidea cancriformis* (94 of the 306 samples), *Rhizoplaca macleanii* (51 samples) and *Lecidella greenii* (37 samples), followed by *Carbonea* sp. 2 (13 samples), *C. vorticosa* (11 samples), *Lecidea polypycnidophora* (10 samples) and *Lecidella siplei* (10 samples; see Supplementary Fig. S5). Nine mycobiont species were found exclusively in area 5 (MDV, 78°S): *Carbonea vorticosa*, *Lecanora* cf. *mons-nivis*, *L.* sp. 2, *Lecidea lapicida*, *L. polypycnidophora*, *L*. sp. 5, *L*. sp. 6*, L*. UCR1 and *Rhizoplaca macleanii.* On the other hand, only the mycobiont species *Lecidea cancriformis* was found in all the six areas; *Lecanora fuscobrunnea* was present in all the areas with the exception of area 2.

The most common photobiont OTUs were *Tr*_A02 (165 of the 281 samples) and *Tr*_S02 (59 samples), both of them occurring in all the six different areas, followed by *Tr*_S18 (32 samples), *Tr*_S15 (10 samples, confined to area 5) and *Tr*_I01 (10 samples). However, of the 149 photobiont samples of area 5, 134 (89,93 %) were assigned to *Tr*_A02. This percentage is much higher than in the other areas (area 1: 44,44 %, area 2: 69,23 %, area 3: 21,74 %, area 4a: 7,69 %, area 4b: 6,67 %), even if those samples with mycobionts occurring exclusively in area 5 (see above) were excluded (76.56 % of the 64 remaining samples are assigned to *Tr*_A02).

The alpha, beta and gamma diversity values are given in Table 2. For the mycobionts, the value of alpha diversity (species richness within a particular area) was the highest in area 5 (8.93) and the lowest in area 4b (1.88). In contrast, for the photobionts, the lowest alpha diversity value was found in area 5 (1.50) and the highest in area 4a (4.06). Thus, referring to this, area 5 plays a remarkable role: compared to the other areas, it shows the highest richness of mycobiont species on the one hand and the lowest richness of photobiont OTUs on the other hand.

**Table 2.** Number of lichen samples, number of identified mycobiont species and photobiont OTUs, as well as alpha, beta and gamma diversity values of mycobiont species/ photobiont OTUs for the different areas.

| Area | Number of lichen samples | Mycobiont species |  |  |  | Photobiont OTUs |  |  |  |
| --- | --- | --- | --- | --- | --- | --- | --- | --- | --- |
|  |  | Number of identified species | Alpha diversity | Beta diversity | Gamma diversity | Number of identified OTUs | Alpha diversity | Beta diversity | Gamma diversity |
| 1 | 28 | 7 | 5.23 | 1.69 | 9.92 | 3 | 2.28 | 1.64 | 3.35 |
| 2 | 13 | 5 | 5.48 |  |  | 3 | 2.20 |  |  |
| 3 | 27 | 7 | 5.99 |  |  | 5 | 3.70 |  |  |
| 4a | 48 | 6 | 3.55 |  |  | 6 | 4.06 |  |  |
| 4b | 31 | 2 | 1.88 |  |  | 4 | 2.36 |  |  |
| 5 | 159 | 16 | 8.93 |  |  | 4 | 1.50 |  |  |

The beta diversity values (extent in changes of species diversity between the areas) for mycobiont species and photobiont OTUs are quite similar (1.69 and 1.64, respectively). This is in contrast to gamma diversity values: the overall diversity for the different areas within the whole region is much higher for mycobiont species (9.92) than for photobiont OTUs (3.35).

### Influence of environmental factors (elevation, precipitation and temperature)

First, the proportion of *Tr*_A02 samples was significantly correlated to BIO10 means of the areas (R = 0.87, p = 0.022; see Supplementary Fig. S6): the higher the temperature mean values of the warmest quarter of an area, the higher the proportion of samples containing photobionts that are assigned to the OTU *Tr*_A02.

The alpha diversity values of mycobiont species significantly positively correlated with BIO10 (R = 0.88, p = 0.021; see Supplementary Fig. S7): the higher the temperature mean values of the warmest quarter, the higher the mycobiont diversity within this particular area.

Furthermore, the differences in mycobiont species community composition were significantly related to BIO10 (constrained principal coordinate analysis: F = 14.7137, p = 0.001, see Supplementary Fig. S8), BIO12 (F = 2.7535, p = 0.012), elevation (F = 2.5108, p = 0.025) and the geographic separation of the samples (Mantel statistic r = 0.1288, p = 0.0002).

The differences in community composition of photobiont OTUs were related significantly to BIO10 (constrained principal coordinate analysis: F = 48.5952, p = 0.001, see Supplementary Fig. S9), BIO12 (F = 4.4848, p = 0.008), elevation (F = 6.8608, p = 0.002), and physical distance (Mantel statistic r = 0.4472, p = 0.0001).

### Haplotype analysis

Haplotype networks were computed for the mycobiont species and photobiont OTUs with h ≥ 2 and at least one haplotype with *n* ≥ 3 (*Carbonea* sp. 2, *Lecanora fuscobrunnea*, *Lecidea cancriformis, Lecidella greenii, L. siplei, L.* sp. nov2 and *Rhizoplaca macleanii,* as well as *Tr*_A02, *Tr*_I01 and *Tr*_S02), in both cases based on nrITS sequence data (Figs. 2 & 3). The samples of *Carbonea vorticosa* (11) were all assigned to a single haplotype, which was also true for *Lecidea polypycnidophora* (10 samples), *Tr*_S15 (10 samples) and *Tr*_S18 (32 samples). Figure 3b and c illustrate the subdivision of *Tr*_I01 (Muggia et al. 2020) into *Tr*_I01j (Leavitt et al. 2015; Ruprecht et al. 2020) and *Tr*_I01k (in this study), and the subdivision of *Tr*_S02 into *Tr*_S02 (Leavitt et al. 2015), and *Tr*_S02b and *Tr*_S02c (Ruprecht et al. 2020).

**Figure 2.**
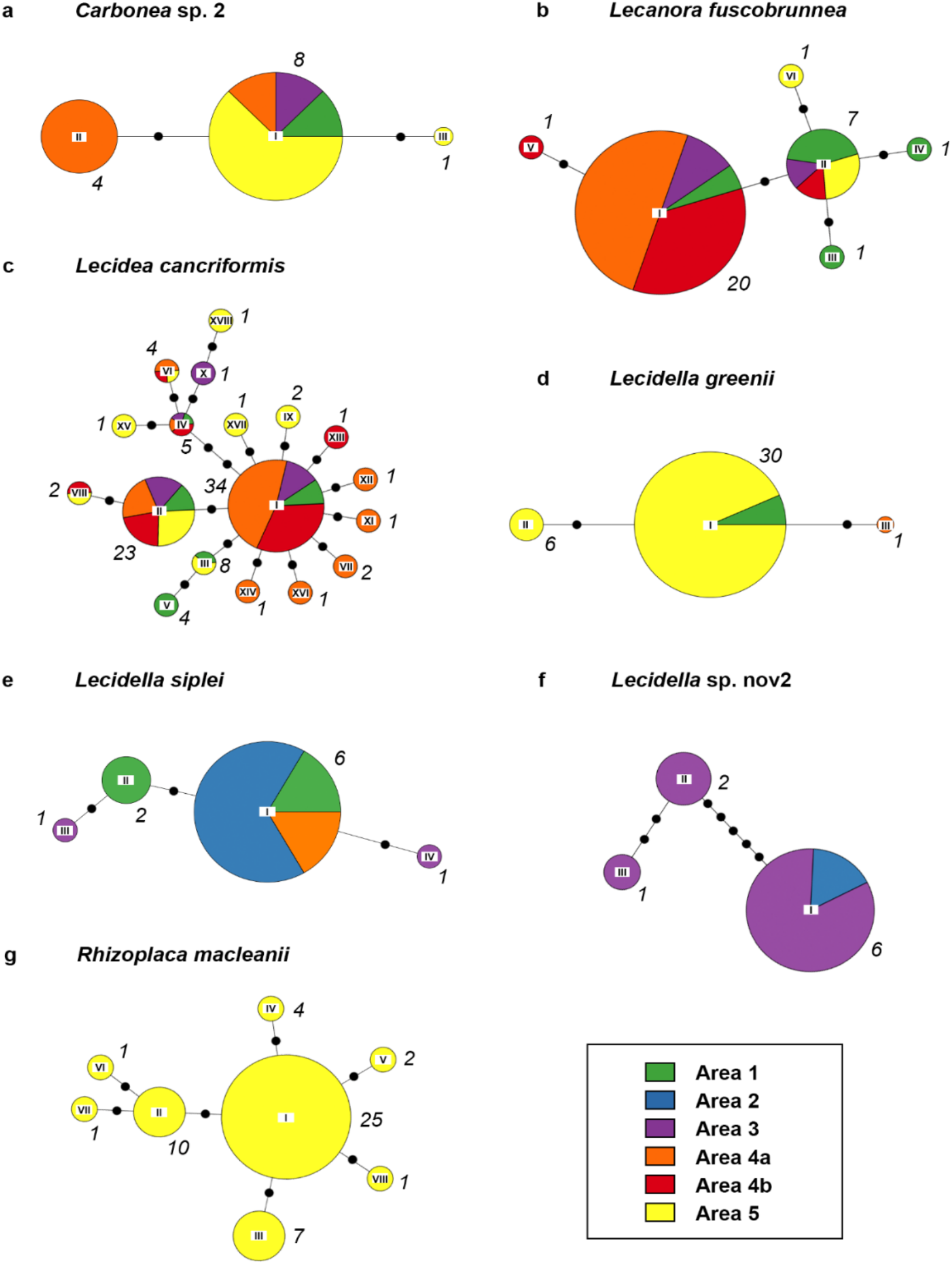
Haplotype networks of mycobiont species with *h* ≥ 2 and at least one haplotype with *n* ≥ 3, showing the spatial distribution within the different areas. Based on nrITS data. (a) *Carbonea* sp. 2, (b) *Lecanora fuscobrunnea,* (c) *Lecidea cancriformis,* (d) *Lecidella greenii,* (e) *Lecidella siplei,* (f) *Lecidella* sp. nov2, (g) *Rhizoplaca macleanii.* Roman numerals at the center of the pie charts refer to the haplotype IDs; the italic numbers next to the pie charts give the total number of samples per haplotype. The circle sizes reflect relative frequency within the species; the frequencies were clustered in ten (e.g. the circles of all haplotypes making up between 20-30 % have the same size). Note: only complete sequences were included.

**Figure 3.**
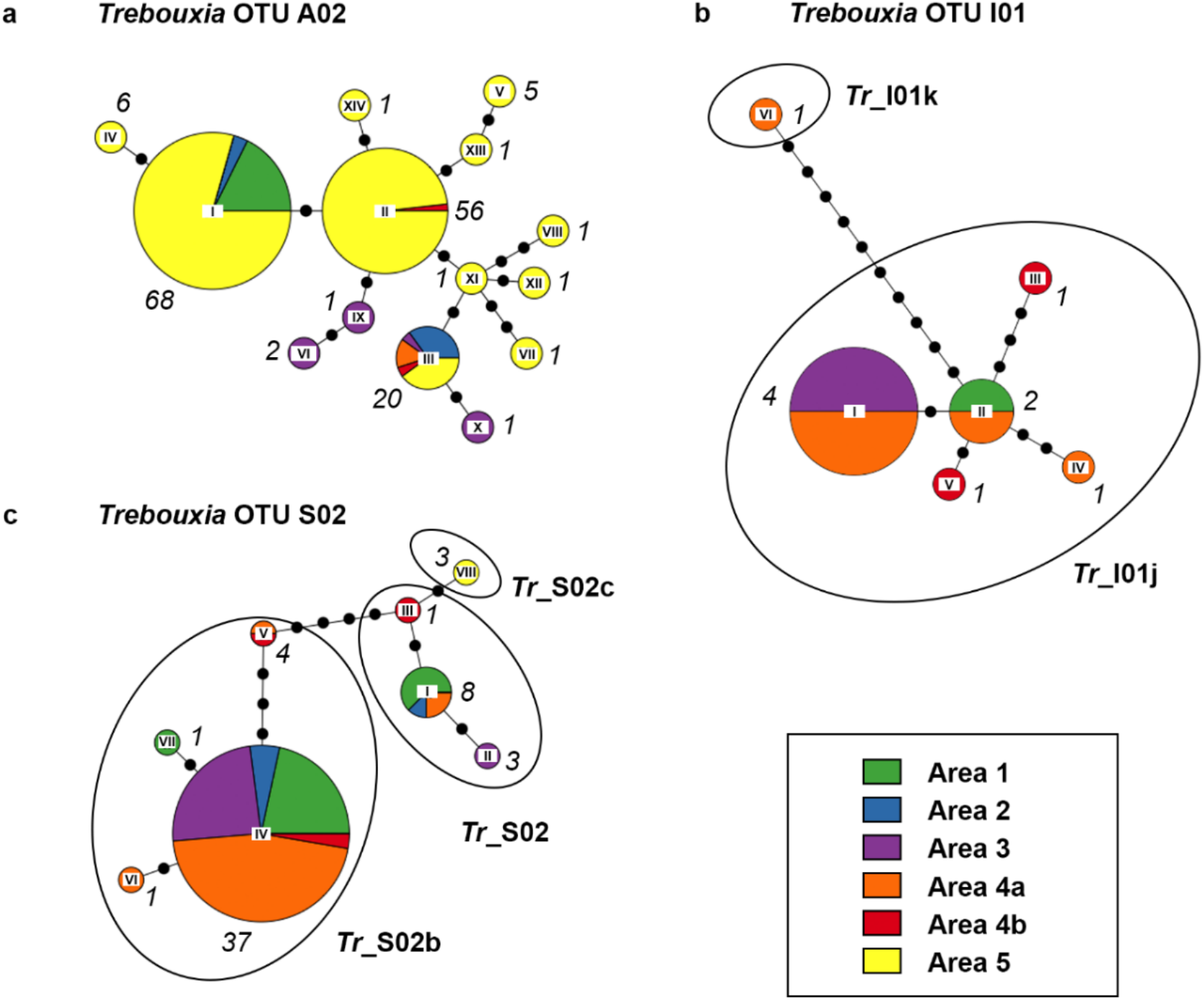
Haplotype networks of photobiont OTUs with *h* ≥ 2 and at least one haplotype with *n* ≥ 3, showing the spatial distribution within the different areas. Based on nrITS data. (a) *Tr*_A02, (b) *Tr*_I01, (c) *Tr*_S02. Roman numerals at the center of the pie charts refer to the haplotype IDs; the italic numbers next to the pie charts give the total number of samples per haplotype. The circle sizes reflect relative frequency within the species; the frequencies were clustered in ten (e.g. the circles of all haplotypes making up between 20-30 % have the same size). Note: only complete sequences were included.

The haplotype networks include pie charts showing the occurrence of the different haplotypes within the different areas. All haplotypes of *Rhizoplaca macleanii* are restricted to area 5, as well as *Lecidella greenii* mainly to area 5 and areas 1 & 4a, and *Lecidella* sp. 2 to areas 2 & 3. However, all other species do not suggest a spatial pattern with different haplotypes being specific for different areas. Moreover, the distribution turned out to be rather unspecific, with a great part of the haplotypes found in multiple areas. For the sake of completeness, additionally, haplotype networks based on multi-locus sequence data were computed for the most abundant mycobiont species and photobiont OTU with multi-locus data available (*Lecidea cancriformis* and *Tr*_S02). Not surprisingly, those networks show a greater number of different haplotypes, but they also do not allow conclusions concerning spatial patterns of area specific haplotypes (see Supplementary Fig. S10).

### Diversity and specificity indices of mycobiont species and photobiont OTUs

The diversity and specificity indices for the different mycobiont species and photobiont OTUs are given in Supplementary Table S6.

For the sample locations of mycobiont species with *n* ≥ 10, BIO10 was strongly correlated to the specificity indices *NRI* (net relatedness index) and significantly correlated to *PSR* (phylogenetic species richness) and *1 – J’* (Pielou evenness index). BIO12 was significantly correlated to *NRI*, *PSR* and *1 – J’*. Figure 4 illustrates these correlations: the higher the BIO10 and BIO12 mean values, the higher the *NRI* (phylogenetic clustering of the photobiont symbiosis partners), the lower the *PSR* (increased phylogenetically relatedness of photobiont symbiosis partners) and the higher *1 – J’* (less numerically evenness of the photobiont symbiosis partners). Thus, for the mean values of the sample locations of a mycobiont species, a comparatively high temperature of the warmest quarter and high annual precipitation occurs with associated photobionts that are phylogenetically clustered and closer related to each other. The lowest values of NRI and the highest values of PSR were developed by *Lecidea cancriformis* and L*ecanora fuscobrunnea*, which also showed the lowest BIO10 and BIO12 mean values at their sample sites. On the contrary, the highest values of NRI and PSR were developed by *Rhizoplaca macleanii*, which also had the highest BIO10 and BIO12 means.

**Figure 4.**
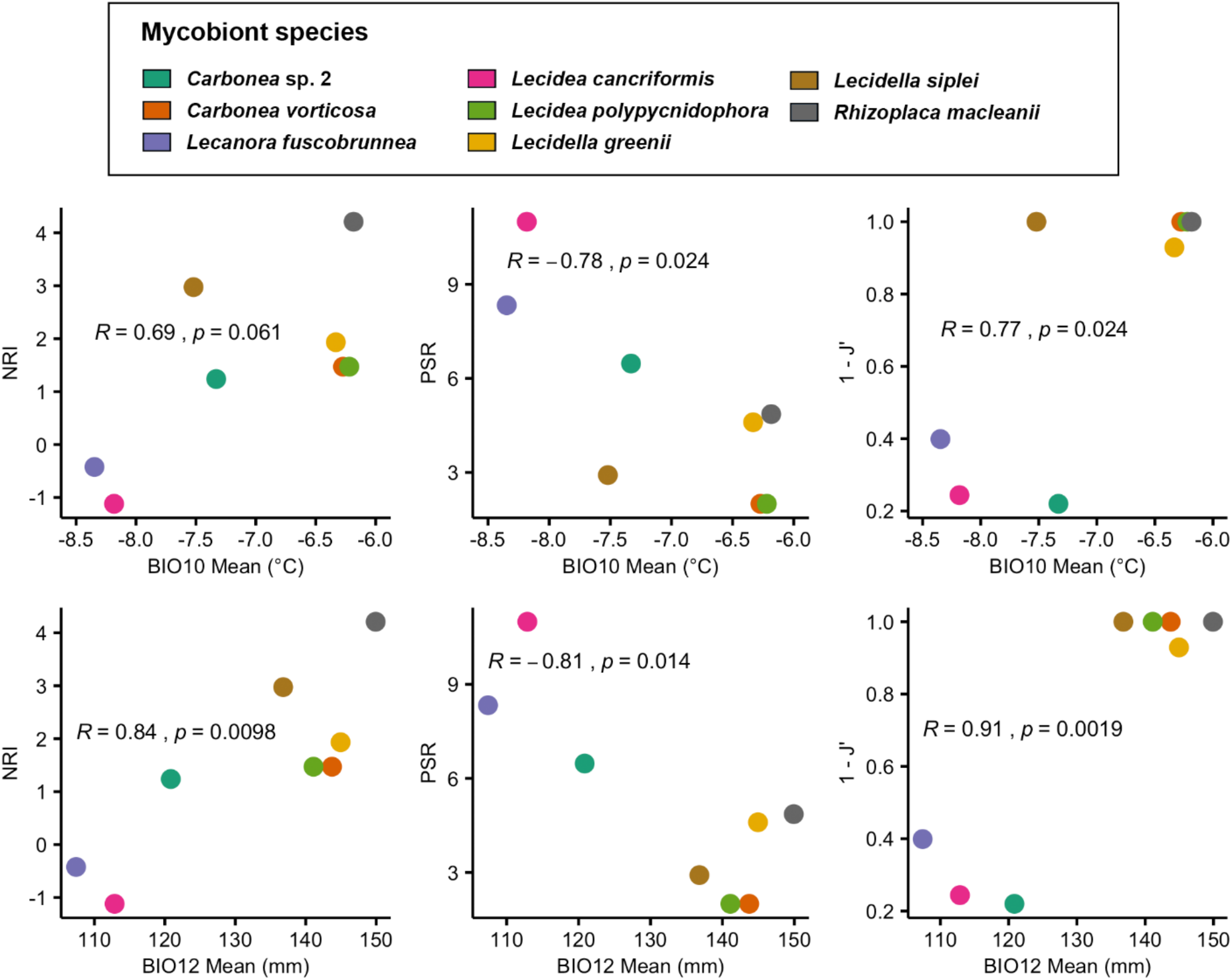
Correlation plots. Specificity indices *NRI* (net relatedness index), *PSR* (phylogenetic species richness and *1 – J’* (Pielou evenness index) against mean values of BIO10 (mean temperature of warmest quarter) and BIO12 (annual precipitation) for mycobiont species with *n* ≥ 10.

For the sample locations of photobiont OTUs with *n* ≥ 10, elevation significantly negatively correlated with *h* (number of haplotypes) and *Hd* (haplotype diversity): the higher the mean elevation of sample sites, the lower the number of haplotypes and the lower the probability that two randomly chosen haplotypes are different (Fig. 5). The highest values of h and *Hd* were for *Tr*_A02, *Tr*_I01 and *Tr*_S02, which occurred at sample sites with comparatively low elevations. In contrast, *Tr*_S15 and *Tr*_S18 occurred at very high elevations and showed very low values of *h* and *Hd*.

**Figure 5.**
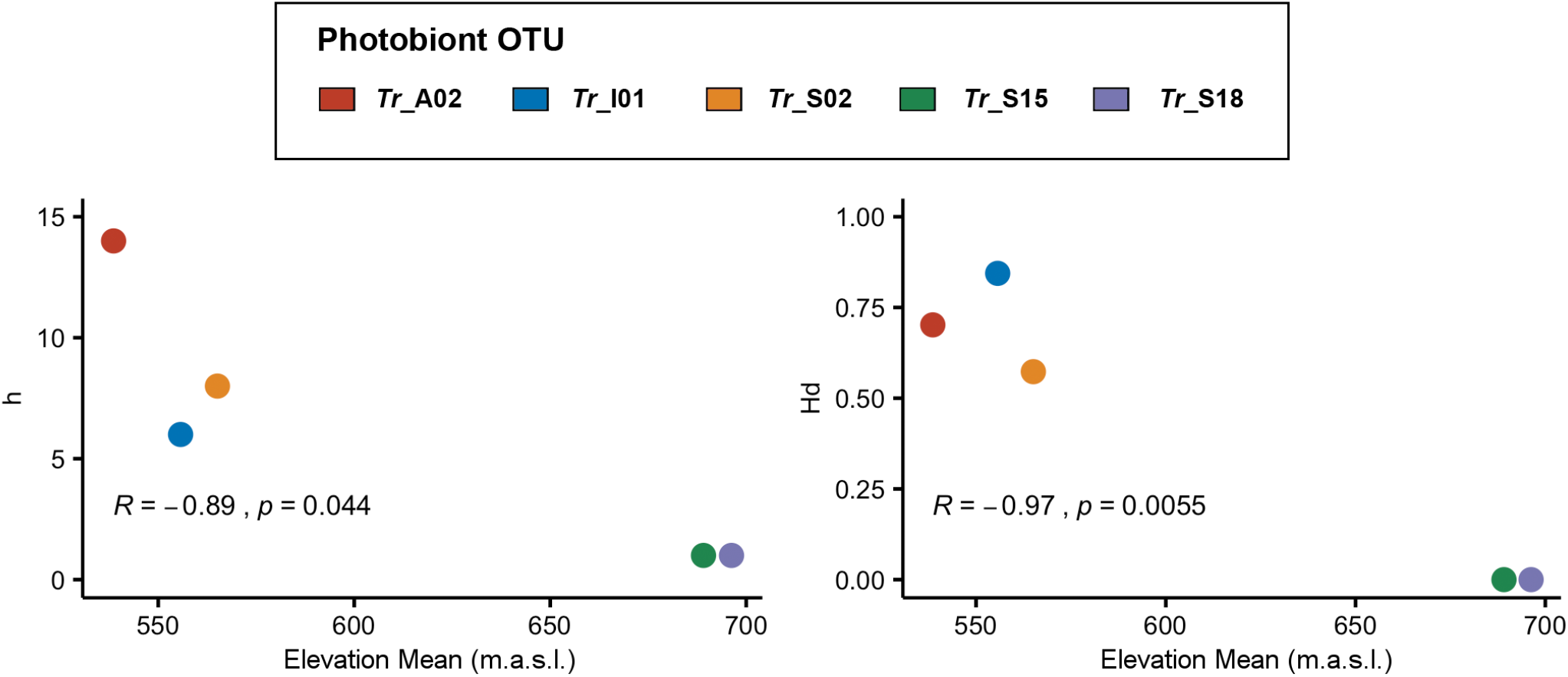
Correlation plots. Diversity indices *h* (number of haplotypes) and *Hd* (haplotype diversity) against mean elevation of sample sites for photobiont OTUs with *n* ≥ 10.

### Analysis of mycobiont-photobiont associations

Bipartite networks were calculated for all associations between mycobiont species (lower level) and the respective photobiont OTUs (higher level) for all areas (Fig. 6). The *H_2_’* value (overall level of complementary specialization of all interacting species) was highest in area 2 (0.921), indicating a network with mostly specialized interactions: within this network, with the exception of *Lecidea andersonii*, the mycobiont species are associated exclusively with one single photobiont OTU. The second highest *H_2_’* value was developed by area 4a (0.710); in contrast, area 4a showed the lowest *H_2_’* value (0.260), with the most abundant mycobiont species *Lecidea cancriformis* showing associations with five different photobiont OTUs. The *H_2_’* values of area 1, area 3 and area 5 indicate medium specification.

**Figure 6.**
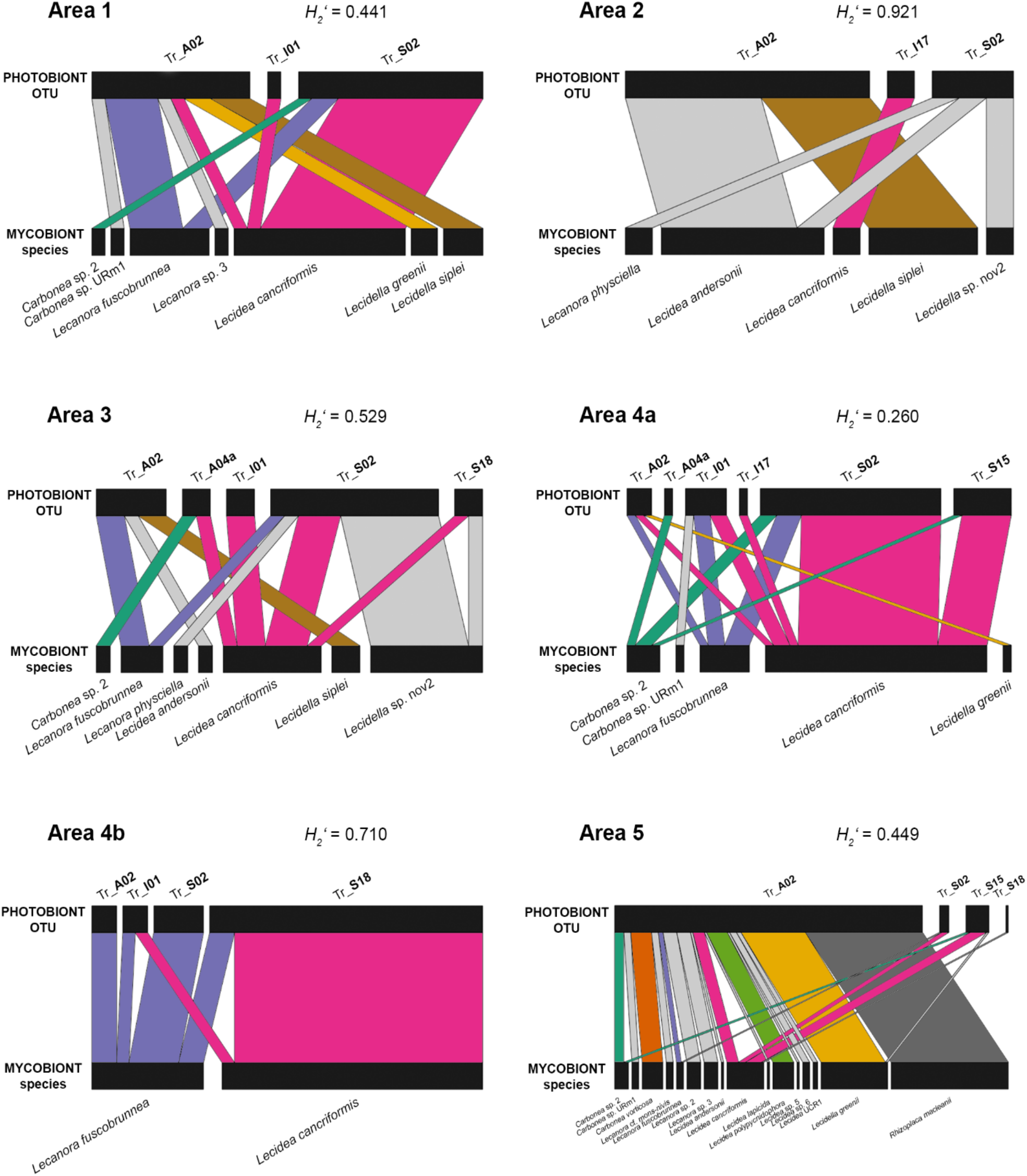
Bipartite networks showing the associations between mycobiont species and photobiont OTUs for the different areas. Rectangles represent species, and the width is proportional to the number of samples. Associated species are linked by lines whose width is proportional to the number of associations.

In addition, the bipartite networks illustrate the different occurrence of mycobiont species and photobiont OTUs within the different areas: For example, in area 1 (and area 2), five (seven) different mycobiont species are associated with only three different photobiont OTUs. In contrast, in area 4b, only two different mycobiont species are associated with four different photobiont OTUs. In area 5, the number of associated photobiont OTUs is also four, but those four OTUs are associated with 16 different mycobiont species.

The network matrix giving all the associations between the mycobiont species and photobiont OTUs is presented in Supplementary Table S7.

## Discussion

The present study investigated the diversity of lecideoid lichens at six different sample areas along a latitudinal gradient (78°S– 85.5°S) along the Transantarctic Mountains, Ross Sea region, at continental Antarctica. The distribution of the different mycobiont species and photobiont OTUs differed considerably between the six sample areas, which is expressed in alpha diversity values (species richness; Supplementary Figure S7). The extreme climate (lowest mean temperatures of the warmest quarter, lowest precipitation) at the areas 4a & b (Darwin Area, ~79.8°S) was reflected in the lowest species richness of mycobionts, and comparatively high species richness of photobiont OTUs. On the other hand, in the climatically mildest area 5 (McMurdo Dry Valleys, 78°S, highest mean temperature, second highest precipitation), the highest species richness of mycobionts and the lowest species richness of photobiont OTUs was found. The number of different photobiont OTUs identified per area is comparatively at a similar level (varying between three and six different groups, Table 2), which is remarkable when considering the great differences in sample sizes (varying between 13 samples (area 2) and 159 samples (area 5), Table 2).

These results are largely consistent with the findings of Colesie et al. (2014), who previously reported that macroclimatic conditions along the latitudinal gradient are not linear. The area at Diamond Hill (79.9°S, part of area 4a, Darwin area, Fig. 1c) addressed in Colesie et al. 2014 showed the lowest species diversity along the latitudinal gradient, which was at least confirmed for area 4b in the current study. Areas 4a and especially 4b are characterized by the harshest climatic conditions such as very low humidity and temperatures, and species numbers in relation to the number of samples are also lowest there; the two generalist species *Lecidea cancriformis* and *Lecanora fuscobrunnea* are dominant. Milder areas, further south (83.5–85.5°S), the McMurdo Dry Valleys (78°S), allow a higher species diversity (Green et al. 2011b; Perez-Ortega et al. 2012). The higher diversity at the southern sites, particularly Mt. Kyffin, appears in part to be due to the occurrence of relic species (Green et al. 2011b). However, Colesie et al. (2014) suggested that physical barriers could be the reason for the low diversity in the Darwin area, but the unspecified haplotype distribution of the widespread species suggest that this is not the case. Since the substrate at all sites is siliceous and there are no other obvious limiting factors, the most likely reason for the limited occurrence of certain species is primarily dependent on abiotic factors, in particular, the environmental conditions caused by geography and macroclimate (Dal Grande et al. 2018).

The uniformity of photobiont OTUs in area 5 is mainly due to a strong dominance (89.93 %) of the OTU *Trebouxia* A02 which occurred in all the six sampling areas. The proportion of photobiont samples assigned to *Tr*_A02 was significantly correlated to the mean value of BIO10 (mean temperature of the warmest month) of the areas. Thus, higher temperatures are related to a higher relative abundance of *Tr*_A02, and colder temperatures to a higher relative abundance of other *Trebouxia* OTUs. This result is in basic agreement with the previous study of Wagner et al. (2020). The community composition of both, mycobiont species as well as photobiont OTUs is significantly related to elevation, BIO10 and BIO12 (annual precipitation). Thus, as sampling sites become more dissimilar in terms of elevation, BIO10 or BIO12, they also become more dissimilar in terms of community composition. These findings are partially supported by Rolshausen et al. (2020) who surveyed *Trebouxia* communities in temperate climates, suggesting that photobionts are replaced by others as environmental conditions change. In addition, significant correlations emerged between the composition of mycobiont and photobiont OTU communities and the geographic separation of the samples: The further the sampling sites are spatially separated, the more dissimilar the corresponding communities become, which is in agreement with Fernandez-Mendoza et al. (2011).

Furthermore, the specificity of mycobiont species towards their photobiont partners was shown to be related to environmental variables; these findings are partially in agreement with the studies of Peksa and Skaloud (2011), Singh et al. (2017) and Rolshausen et al. (2018), who reported climate as well as substrate a selective pressure in terms of increased specificity of mycobiont-photobiont interactions. However, the current study has shown that a higher value of BIO10 correlated with a higher phylogenetic clustering of the symbiotic partners of a single mycobiont species (higher *1 – J’* and *NRI* values) and a closer phylogenetic relatedness of these photobionts (lower PSR values). Similarly, the specificity of mycobiont species towards their photobiont symbiosis partners also correlated with BIO12 mean values: A higher value of BIO12 is related to higher values of *1 – J’* and *NRI* and to lower values of *PSR*. Consequently, the mycobiont species with *n* ≥ 10 showing the highest BIO10 and BIO12 mean values at its sample locations (*Rhizoplaca macleanii*) also had the highest value of *NRI* and a rather low value of *PSR*, as it was solely associated with *Tr*_A02. On the other hand, the two mycobiont species with *n* ≥ 10 showing the lowest mean values of BIO10 and BIO12 (*Lecanora fuscobrunnea* and *Lecidea cancriformis)* exhibited the lowest values of *NRI* and the highest values of *PSR*, as they had associations with the phylogenetically distinct *Trebouxia* OTUs A02, I01, S02 and S18 (*Lecanora fuscobrunnea*) or all seven *Trebouxia* OTUs of this study (*Lecidea cancriformis*), respectively. Additionally, *Lecanora fuscobrunnea* and *Lecidea cancriformis* were the two most widespread species that occurred in five of the six (*L. fuscobrunnea*) or all the six different areas (*L. cancriformis*). This result is in agreement with former studies that had shown that *L. cancriformis* is able to associate with all known photobiont species, and is one of the most widespread lichens in continental Antarctica (Castello 2003; Ruprecht et al. 2012a; Ruprecht et al. 2010; Wagner et al. 2020). Previous studies suggested that a higher photobiont diversity within a single lichen species is indicative of a lower selectivity by the mycobiont, and that this condition is related to enhanced colonization ability (Blaha et al. 2006; Guzow-Krzeminska 2006; Wirtz et al. 2003). According to a model developed by Yahr et al. (2006), selectivity may vary between habitats and may enable lichens to select a photobiont that is well adapted to conditions of the local environmental. These photobiont switches were suggested to increase the geographical range and ecological niche of lichen mycobionts, but may also lead to genetic isolation between mycobiont populations and thus drive their evolution (Fernandez-Mendoza et al. 2011). More generally, flexibility concerning the partner choice has been considered to be an adaptive strategy to survive harsher environmental conditions (Engelen et al. 2016; Leavitt et al. 2013; Singh et al. 2017; Werth and Sork 2010).

Dal Grande et al. (2018) reported elevational preferences for some *Trebouxia* taxa at the OTU level at a mountain range in central Spain, covering an elevational gradient of 1400 m. Additionally, in the present study, the mean elevation of photobiont OTUs were negatively correlated to differences in diversity indices: the dominant photobiont OTU *Tr*_A02, occurring in all the six different areas, exhibited the lowest mean value of elevation of sample sites and had the highest number of haplotypes and the highest value of haplotype diversity. On the other hand, the OTUs *Tr*_S15 and *Tr*_S18 had the highest mean elevations and the lowest values of *h* and *Hd*. Thus, higher mean elevation of photobiont OTUs were significantly related to a lower number of haplotypes (*h*) and a lower haplotype diversity (*Hd*).

## Conclusions

Lichens and their myco-/photobiont associations clearly show environmental preferences and therefore are useful as bioindicators. The *Trebouxia* OTU A02 occurred in all the six different areas and was dominant in milder areas, whereas in colder areas, a higher relative abundance of other *Trebouxia* OTUs was found. Accordingly, mycobiont species occurring in milder areas (like *Carbonea vorticosa*, *Lecidella greenii* and *Rhizoplaca macleanii*) are almost exclusively associated with *Tr*_A02, while the generalist mycobiont species *Lecidea cancriformis* und *Lecanora fuscobrunnea*, occurring in a broad range of climatically different environments, show associations with phylogenetically distinct photobiont OTUs. However, if they are the only lecideoid lichen species present in certain areas, then they are also meaningful bioindicators of extreme climatic conditions.

## Acknowledgements

We want to thank D. Laina, G. Zimmermann, the IDA Lab (Salzburg, A) and T.G.A. Green (Waikato, NZ) for various help and valuable advice and suggestions.

## Additional information

### Supplementary information

Supplementary Material 1: Tables, Supplementary Material 2: Figures

## Supplementary Material 1: Tables

**Supplementary Table S1.**
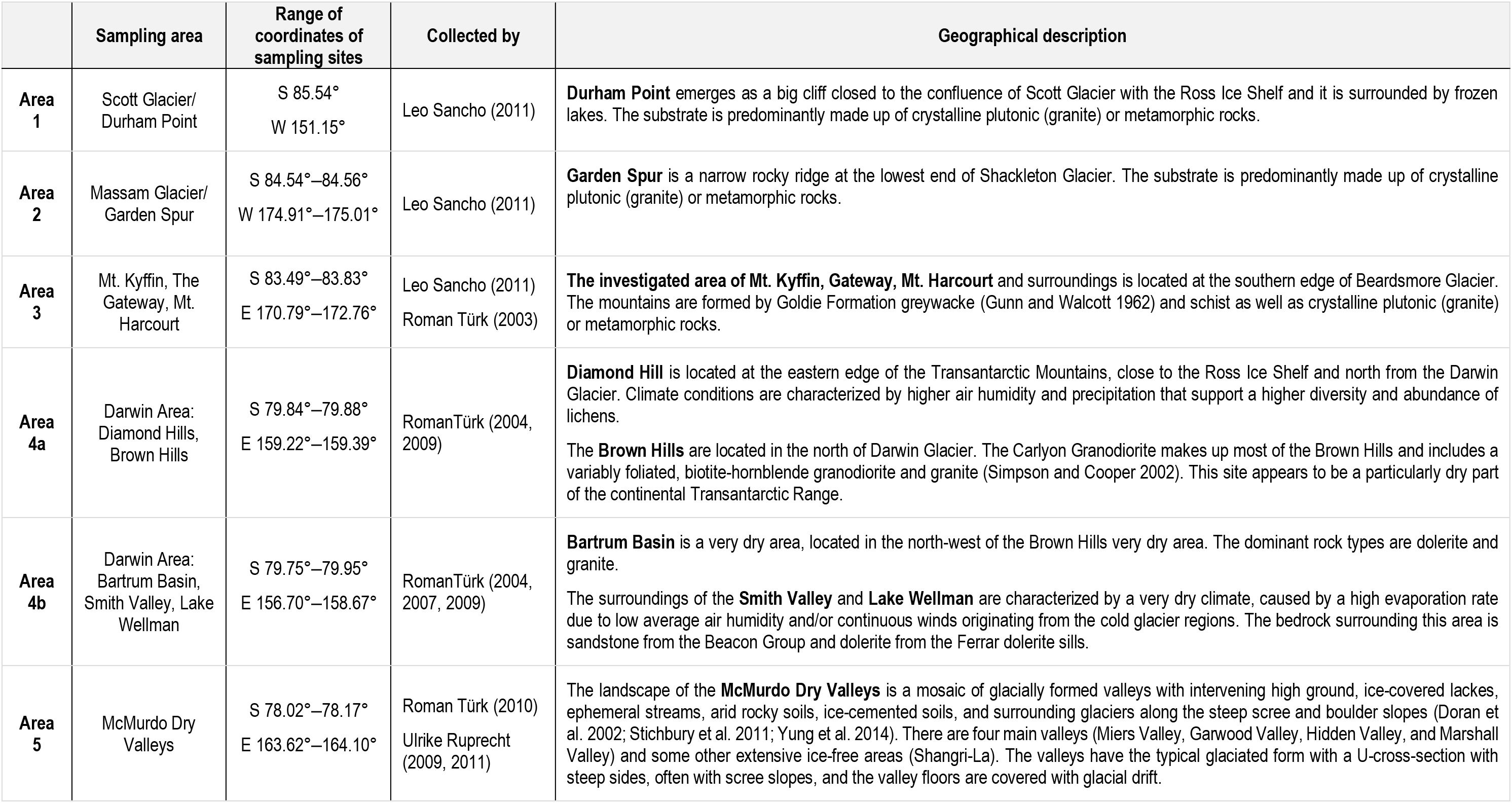
Geographical description of sampling sites.

|  | Sampling area | Range of coordinates of sampling sites | Collected by | Geographical description |
| --- | --- | --- | --- | --- |
| **Area 1** | Scott Glacier/<br>Durham Point | S 85.54°<br>W 151.15° | Leo Sancho (2011) | **Durham Point** emerges as a big cliff closed to the confluence of Scott Glacier with the Ross Ice Shelf and it is surrounded by frozen lakes. The substrate is predominantly made up of crystalline plutonic (granite) or metamorphic rocks. |
| **Area 2** | Massam Glacier/<br>Garden Spur | S 84.54°–84.56°<br>W 174.91°–175.01° | Leo Sancho (2011) | **Garden Spur** is a narrow rocky ridge at the lowest end of Shackleton Glacier. The substrate is predominantly made up of crystalline plutonic (granite) or metamorphic rocks. |
| **Area 3** | Mt. Kyffin, The<br>Gateway, Mt.<br>Harcourt | S 83.49°–83.83°<br>E 170.79°–172.76° | Leo Sancho (2011)<br>Roman Türk (2003) | **The investigated area of Mt. Kyffin, Gateway, Mt. Harcourt** and surroundings is located at the southern edge of Beardsmore Glacier. The mountains are formed by Goldie Formation greywacke (Gunn and Walcott 1962) and schist as well as crystalline plutonic (granite) or metamorphic rocks. |
| **Area 4a** | Darwin Area:<br>Diamond Hills,<br>Brown Hills | S 79.84°–79.88°<br>E 159.22°–159.39° | RomanTürk (2004,<br>2009) | **Diamond Hill** is located at the eastern edge of the Transantarctic Mountains, close to the Ross Ice Shelf and north from the Darwin Glacier. Climate conditions are characterized by higher air humidity and precipitation that support a higher diversity and abundance of lichens.<br>The **Brown Hills** are located in the north of Darwin Glacier. The Carlyon Granodiorite makes up most of the Brown Hills and includes a variably foliated, biotite-hornblende granodiorite and granite (Simpson and Cooper 2002). This site appears to be a particularly dry part of the continental Transantarctic Range. |
| **Area 4b** | Darwin Area:<br>Bartrum Basin,<br>Smith Valley, Lake<br>Wellman | S 79.75°–79.95°<br>E 156.70°–158.67° | RomanTürk (2004,<br>2007, 2009) | **Bartrum Basin** is a very dry area, located in the north-west of the Brown Hills very dry area. The dominant rock types are dolerite and granite.<br>The surroundings of the **Smith Valley** and **Lake Wellman** are characterized by a very dry climate, caused by a high evaporation rate due to low average air humidity and/or continuous winds originating from the cold glacier regions. The bedrock surrounding this area is sandstone from the Beacon Group and dolerite from the Ferrar dolerite sills. |
| **Area 5** | McMurdo Dry<br>Valleys | S 78.02°–78.17°<br>E 163.62°–164.10° | Roman Türk (2010)<br>Ulrike Ruprecht<br>(2009, 2011) | The landscape of the **McMurdo Dry Valleys** is a mosaic of glacially formed valleys with intervening high ground, ice-covered lakes, ephemeral streams, arid rocky soils, ice-cemented soils, and surrounding glaciers along the steep scree and boulder slopes (Doran et al. 2002; Stichbury et al. 2011; Yung et al. 2014). There are four main valleys (Miers Valley, Garwood Valley, Hidden Valley, and Marshall Valley) and some other extensive ice-free areas (Shangri-La). The valleys have the typical glaciated form with a U-cross-section with steep sides, often with scree slopes, and the valley floors are covered with glacial drift. |

**Supplementary Table S2.**
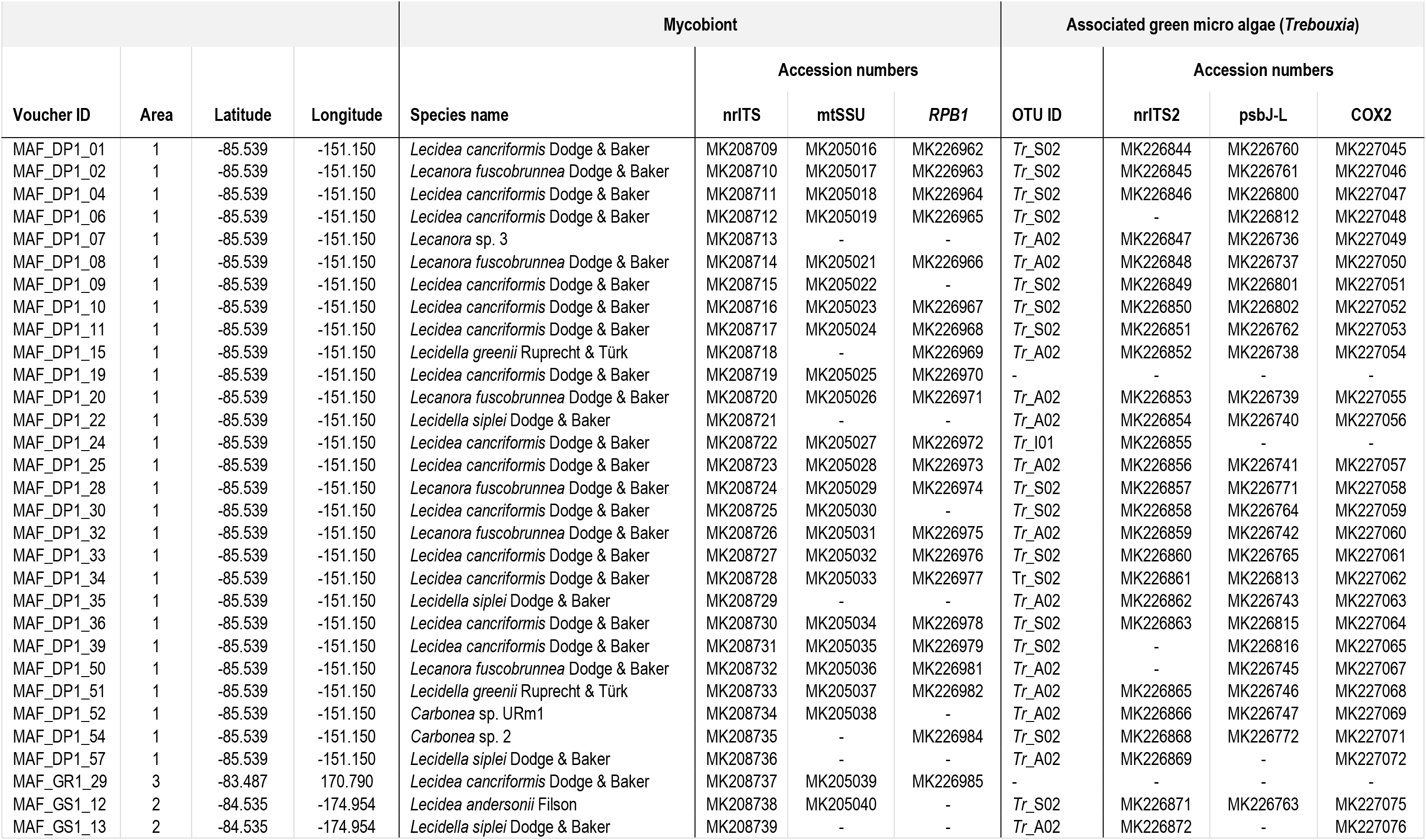
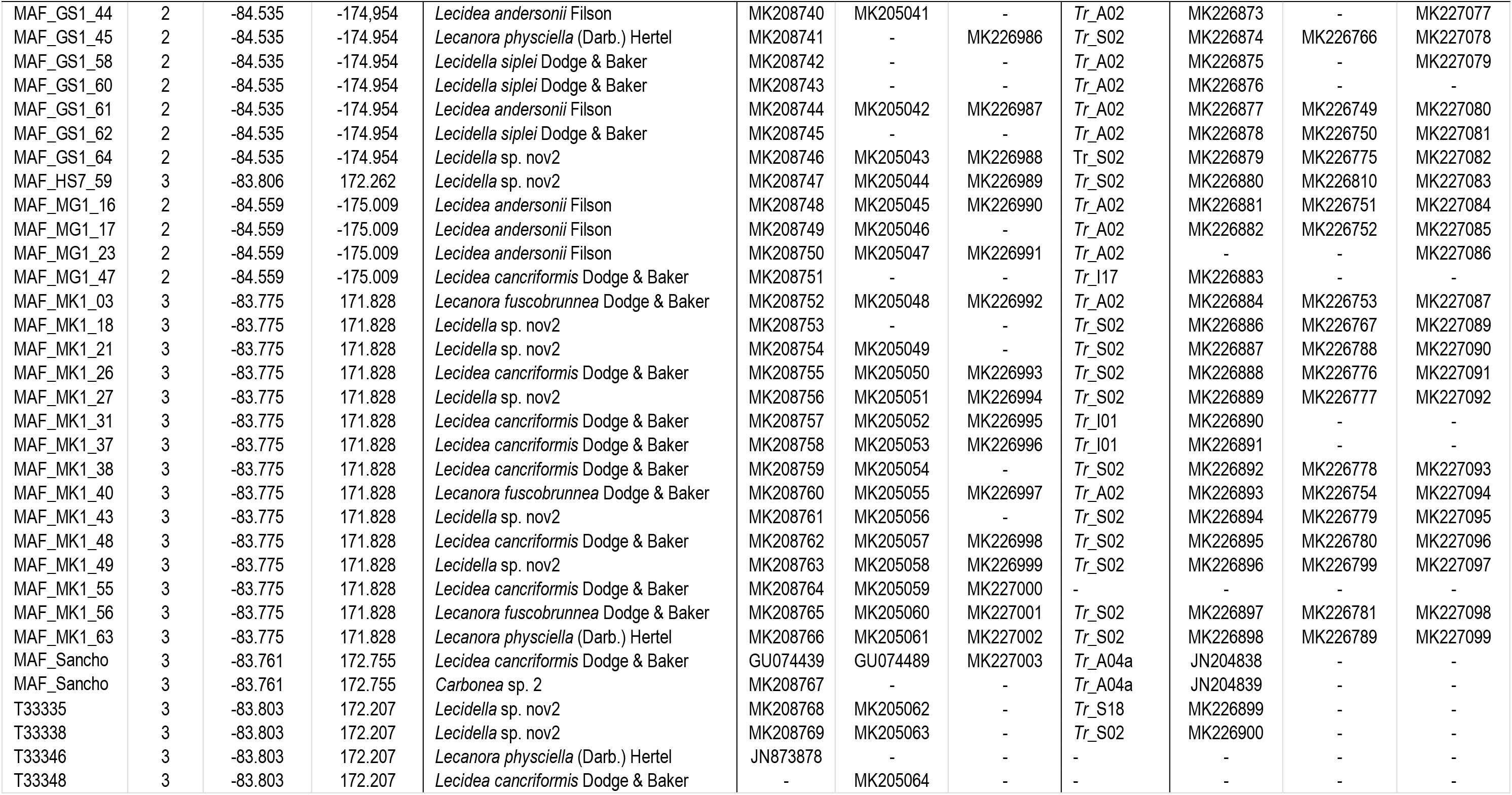
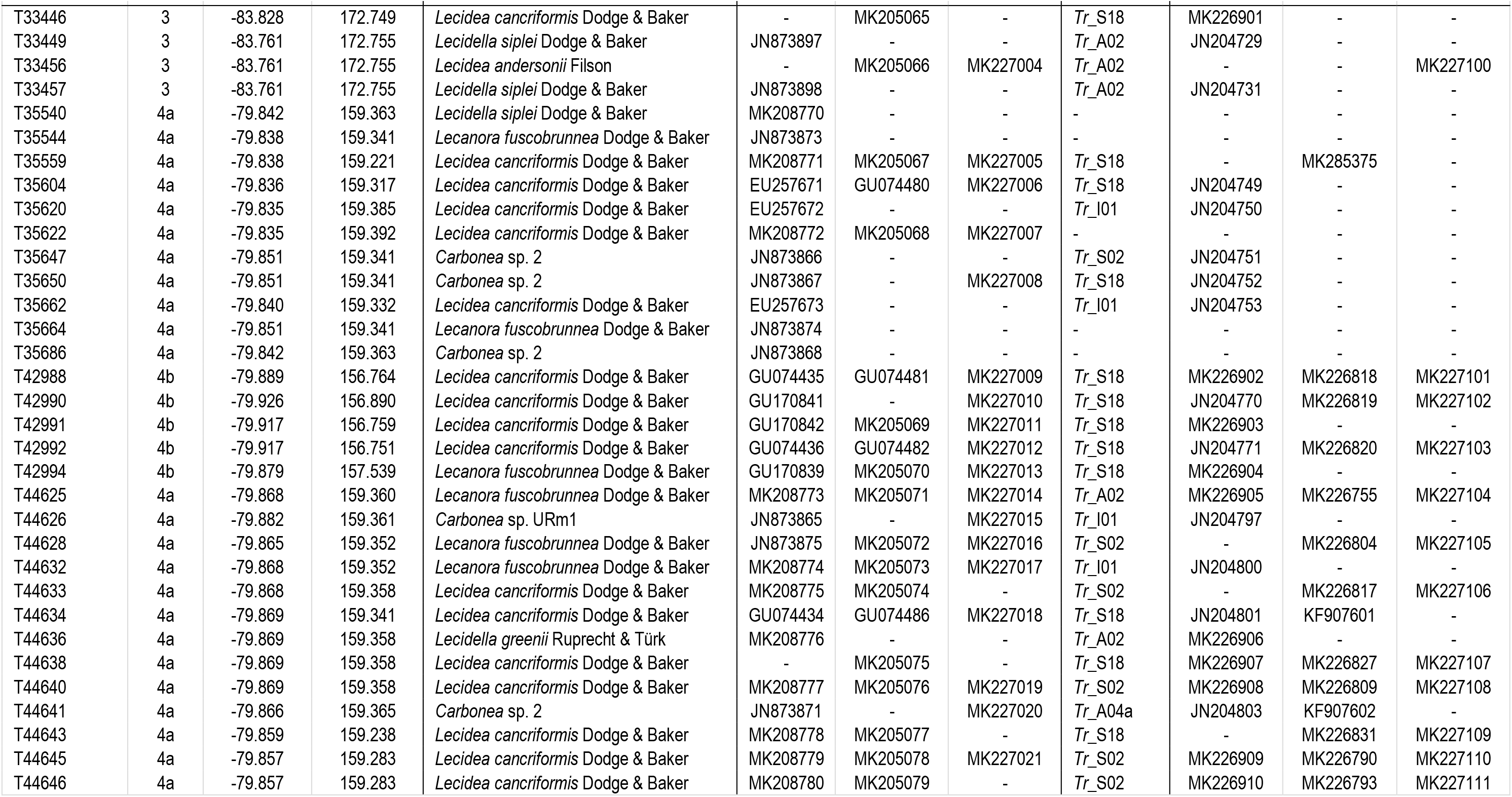
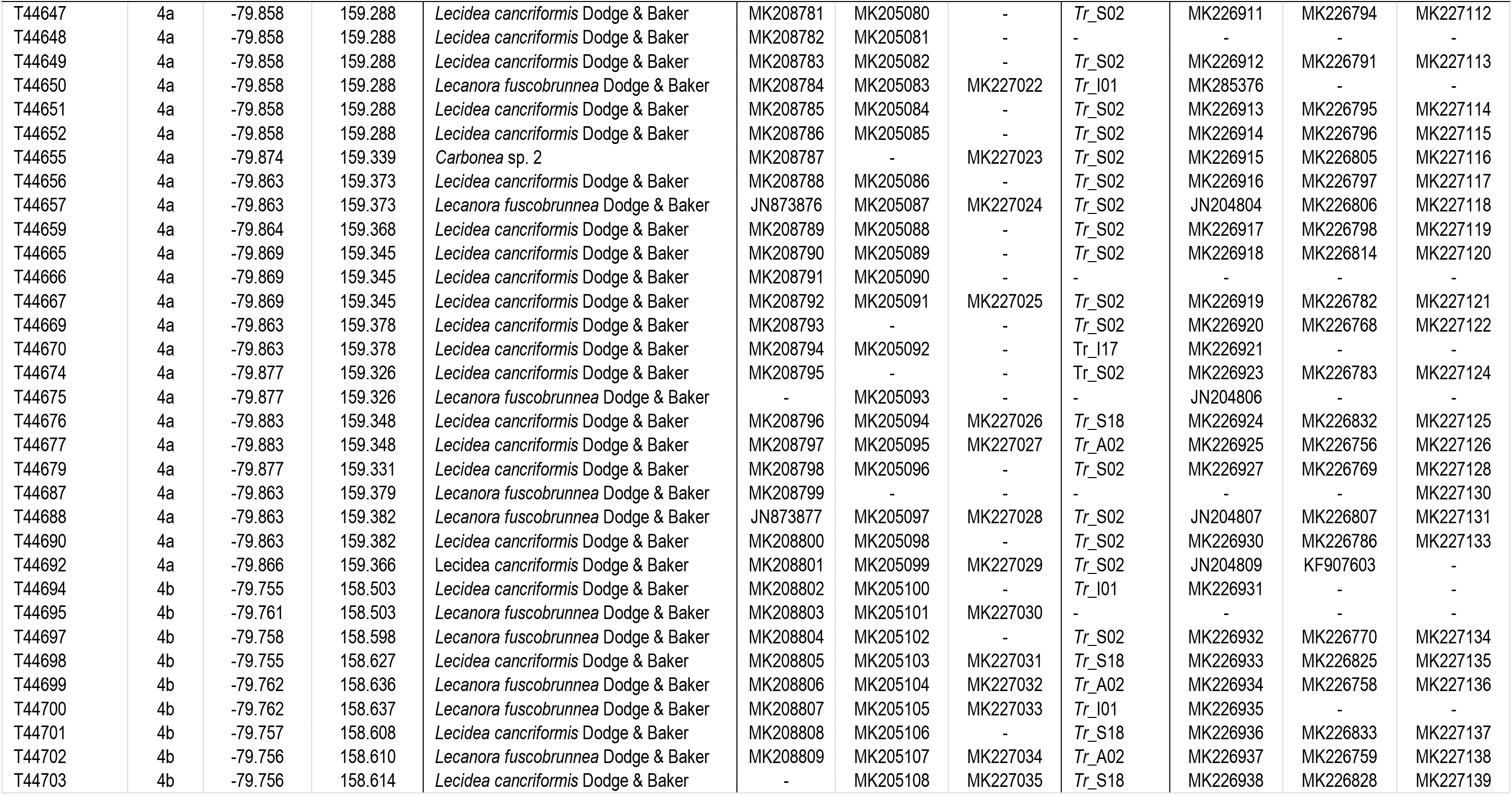
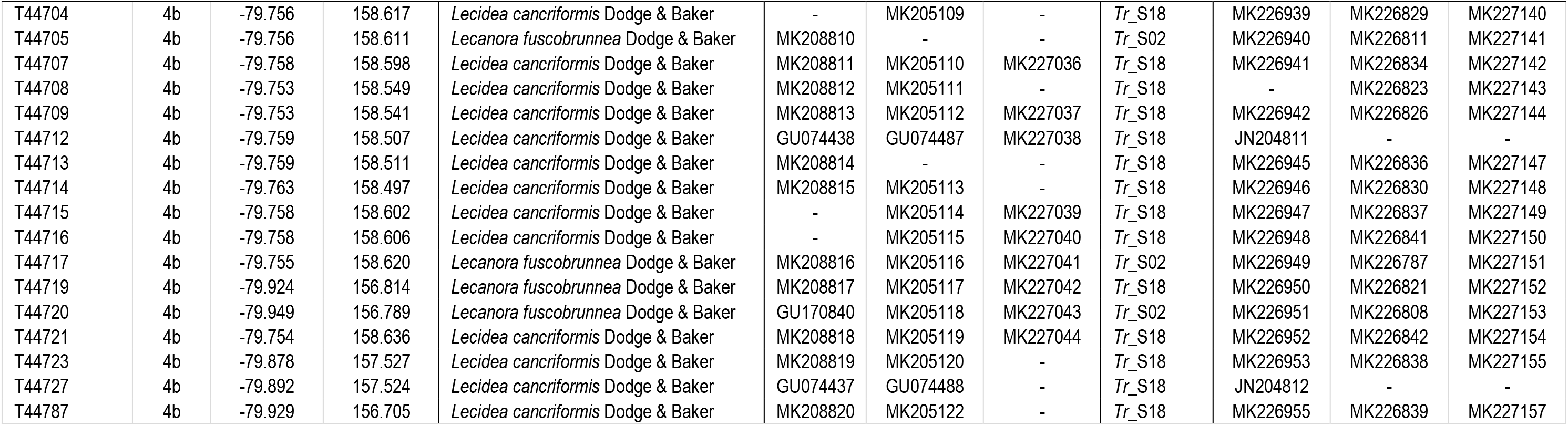
Samples used in this study, with information on collecting localities and Genbank accession numbers of different markers.

| Voucher ID | Area | Latitude | Longitude | Mycobiont |  |  | Associated green micro algae (*Trebouxia*) |  |  |  |  |
| --- | --- | --- | --- | --- | --- | --- | --- | --- | --- | --- | --- |
|  |  |  |  | Species name | Accession numbers |  |  | OTU ID | Accession numbers |  |  |
|  |  |  |  |  | nrITS | mtSSU | RPB1 |  | nrITS2 | psbJ-L | COX2 |
| MAF_DP1_01 | 1 | -85.539 | -151.150 | *Lecidea cancriiformis* Dodge & Baker | MK208709 | MK205016 | MK226962 | *Tr_S02* | MK226844 | MK226760 | MK227045 |
| MAF_DP1_02 | 1 | -85.539 | -151.150 | *Lecanora fuscobrunnea* Dodge & Baker | MK208710 | MK205017 | MK226963 | *Tr_S02* | MK226845 | MK226761 | MK227046 |
| MAF_DP1_04 | 1 | -85.539 | -151.150 | *Lecidea cancriiformis* Dodge & Baker | MK208711 | MK205018 | MK226964 | *Tr_S02* | MK226846 | MK226800 | MK227047 |
| MAF_DP1_06 | 1 | -85.539 | -151.150 | *Lecidea cancriiformis* Dodge & Baker | MK208712 | MK205019 | MK226965 | *Tr_S02* | - | MK226812 | MK227048 |
| MAF_DP1_07 | 1 | -85.539 | -151.150 | *Lecanora* sp. 3 | MK208713 | - | - | *Tr_A02* | MK226847 | MK226736 | MK227049 |
| MAF_DP1_08 | 1 | -85.539 | -151.150 | *Lecanora fuscobrunnea* Dodge & Baker | MK208714 | MK205021 | MK226966 | *Tr_A02* | MK226848 | MK226737 | MK227050 |
| MAF_DP1_09 | 1 | -85.539 | -151.150 | *Lecidea cancriiformis* Dodge & Baker | MK208715 | MK205022 | - | *Tr_S02* | MK226849 | MK226801 | MK227051 |
| MAF_DP1_10 | 1 | -85.539 | -151.150 | *Lecidea cancriiformis* Dodge & Baker | MK208716 | MK205023 | MK226967 | *Tr_S02* | MK226850 | MK226802 | MK227052 |
| MAF_DP1_11 | 1 | -85.539 | -151.150 | *Lecidea cancriiformis* Dodge & Baker | MK208717 | MK205024 | MK226968 | *Tr_S02* | MK226851 | MK226762 | MK227053 |
| MAF_DP1_15 | 1 | -85.539 | -151.150 | *Lecidella greenii* Ruprecht & Türk | MK208718 | - | MK226969 | *Tr_A02* | MK226852 | MK226738 | MK227054 |
| MAF_DP1_19 | 1 | -85.539 | -151.150 | *Lecidea cancriiformis* Dodge & Baker | MK208719 | MK205025 | MK226970 | - | - | - | - |
| MAF_DP1_20 | 1 | -85.539 | -151.150 | *Lecanora fuscobrunnea* Dodge & Baker | MK208720 | MK205026 | MK226971 | *Tr_A02* | MK226853 | MK226739 | MK227055 |
| MAF_DP1_22 | 1 | -85.539 | -151.150 | *Lecidella siplei* Dodge & Baker | MK208721 | - | - | *Tr_A02* | MK226854 | MK226740 | MK227056 |
| MAF_DP1_24 | 1 | -85.539 | -151.150 | *Lecidea cancriiformis* Dodge & Baker | MK208722 | MK205027 | MK226972 | *Tr_I01* | MK226855 | - | - |
| MAF_DP1_25 | 1 | -85.539 | -151.150 | *Lecidea cancriiformis* Dodge & Baker | MK208723 | MK205028 | MK226973 | *Tr_A02* | MK226856 | MK226741 | MK227057 |
| MAF_DP1_28 | 1 | -85.539 | -151.150 | *Lecanora fuscobrunnea* Dodge & Baker | MK208724 | MK205029 | MK226974 | *Tr_S02* | MK226857 | MK226771 | MK227058 |
| MAF_DP1_30 | 1 | -85.539 | -151.150 | *Lecidea cancriiformis* Dodge & Baker | MK208725 | MK205030 | - | *Tr_S02* | MK226858 | MK226764 | MK227059 |
| MAF_DP1_32 | 1 | -85.539 | -151.150 | *Lecanora fuscobrunnea* Dodge & Baker | MK208726 | MK205031 | MK226975 | *Tr_A02* | MK226859 | MK226742 | MK227060 |
| MAF_DP1_33 | 1 | -85.539 | -151.150 | *Lecidea cancriiformis* Dodge & Baker | MK208727 | MK205032 | MK226976 | *Tr_S02* | MK226860 | MK226765 | MK227061 |
| MAF_DP1_34 | 1 | -85.539 | -151.150 | *Lecidea cancriiformis* Dodge & Baker | MK208728 | MK205033 | MK226977 | *Tr_S02* | MK226861 | MK226813 | MK227062 |
| MAF_DP1_35 | 1 | -85.539 | -151.150 | *Lecidella siplei* Dodge & Baker | MK208729 | - | - | *Tr_A02* | MK226862 | MK226743 | MK227063 |
| MAF_DP1_36 | 1 | -85.539 | -151.150 | *Lecidea cancriiformis* Dodge & Baker | MK208730 | MK205034 | MK226978 | *Tr_S02* | MK226863 | MK226815 | MK227064 |
| MAF_DP1_39 | 1 | -85.539 | -151.150 | *Lecidea cancriiformis* Dodge & Baker | MK208731 | MK205035 | MK226979 | *Tr_S02* | - | MK226816 | MK227065 |
| MAF_DP1_50 | 1 | -85.539 | -151.150 | *Lecanora fuscobrunnea* Dodge & Baker | MK208732 | MK205036 | MK226981 | *Tr_A02* | - | MK226745 | MK227067 |
| MAF_DP1_51 | 1 | -85.539 | -151.150 | *Lecidella greenii* Ruprecht & Türk | MK208733 | MK205037 | MK226982 | *Tr_A02* | MK226865 | MK226746 | MK227068 |
| MAF_DP1_52 | 1 | -85.539 | -151.150 | *Carbonea* sp. URm1 | MK208734 | MK205038 | - | *Tr_A02* | MK226866 | MK226747 | MK227069 |
| MAF_DP1_54 | 1 | -85.539 | -151.150 | *Carbonea* sp. 2 | MK208735 | - | MK226984 | *Tr_S02* | MK226868 | MK226772 | MK227071 |
| MAF_DP1_57 | 1 | -85.539 | -151.150 | *Lecidella siplei* Dodge & Baker | MK208736 | - | - | *Tr_A02* | MK226869 | - | MK227072 |
| MAF_GR1_29 | 3 | -83.487 | 170.790 | *Lecidea cancriiformis* Dodge & Baker | MK208737 | MK205039 | MK226985 | - | - | - | - |
| MAF_GS1_12 | 2 | -84.535 | -174.954 | *Lecidea andersonii* Filson | MK208738 | MK205040 | - | *Tr_S02* | MK226871 | MK226763 | MK227075 |
| MAF_GS1_13 | 2 | -84.535 | -174.954 | *Lecidella siplei* Dodge & Baker | MK208739 | - | - | *Tr_A02* | MK226872 | - | MK227076 |

|  |  |  |  | Mycobiont |  |  | Associated green micro algae (*Trebouxia*) |  |  |  |  |
| --- | --- | --- | --- | --- | --- | --- | --- | --- | --- | --- | --- |
| Voucher ID | Area | Latitude | Longitude | Species name | Accession numbers |  |  | OTU ID | Accession numbers |  |  |
|  |  |  |  |  | nrITS | mtSSU | RPB1 |  | nrITS2 | psbJ-L | COX2 |
| MAF_GS1_44 | 2 | -84.535 | -174.954 | *Lecidea andersonii* Filson | MK208740 | MK205041 | - | *Tr_A02* | MK226873 | - | MK227077 |
| MAF_GS1_45 | 2 | -84.535 | -174.954 | *Lecanora physciella* (Darb.) Hertel | MK208741 | - | MK226986 | *Tr_S02* | MK226874 | MK226766 | MK227078 |
| MAF_GS1_58 | 2 | -84.535 | -174.954 | *Lecidella siplei* Dodge & Baker | MK208742 | - | - | *Tr_A02* | MK226875 | - | MK227079 |
| MAF_GS1_60 | 2 | -84.535 | -174.954 | *Lecidella siplei* Dodge & Baker | MK208743 | - | - | *Tr_A02* | MK226876 | - | - |
| MAF_GS1_61 | 2 | -84.535 | -174.954 | *Lecidea andersonii* Filson | MK208744 | MK205042 | MK226987 | *Tr_A02* | MK226877 | MK226749 | MK227080 |
| MAF_GS1_62 | 2 | -84.535 | -174.954 | *Lecidella siplei* Dodge & Baker | MK208745 | - | - | *Tr_A02* | MK226878 | MK226750 | MK227081 |
| MAF_GS1_64 | 2 | -84.535 | -174.954 | *Lecidella* sp. nov2 | MK208746 | MK205043 | MK226988 | *Tr_S02* | MK226879 | MK226775 | MK227082 |
| MAF_HS7_59 | 3 | -83.806 | 172.262 | *Lecidella* sp. nov2 | MK208747 | MK205044 | MK226989 | *Tr_S02* | MK226880 | MK226810 | MK227083 |
| MAF_MG1_16 | 2 | -84.559 | -175.009 | *Lecidea andersonii* Filson | MK208748 | MK205045 | MK226990 | *Tr_A02* | MK226881 | MK226751 | MK227084 |
| MAF_MG1_17 | 2 | -84.559 | -175.009 | *Lecidea andersonii* Filson | MK208749 | MK205046 | - | *Tr_A02* | MK226882 | MK226752 | MK227085 |
| MAF_MG1_23 | 2 | -84.559 | -175.009 | *Lecidea andersonii* Filson | MK208750 | MK205047 | MK226991 | *Tr_A02* | - | - | MK227086 |
| MAF_MG1_47 | 2 | -84.559 | -175.009 | *Lecidea cancriformis* Dodge & Baker | MK208751 | - | - | *Tr_I17* | MK226883 | - | - |
| MAF_MK1_03 | 3 | -83.775 | 171.828 | *Lecanora fuscobrunnea* Dodge & Baker | MK208752 | MK205048 | MK226992 | *Tr_A02* | MK226884 | MK226753 | MK227087 |
| MAF_MK1_18 | 3 | -83.775 | 171.828 | *Lecidella* sp. nov2 | MK208753 | - | - | *Tr_S02* | MK226886 | MK226767 | MK227089 |
| MAF_MK1_21 | 3 | -83.775 | 171.828 | *Lecidella* sp. nov2 | MK208754 | MK205049 | - | *Tr_S02* | MK226887 | MK226788 | MK227090 |
| MAF_MK1_26 | 3 | -83.775 | 171.828 | *Lecidea cancriformis* Dodge & Baker | MK208755 | MK205050 | MK226993 | *Tr_S02* | MK226888 | MK226776 | MK227091 |
| MAF_MK1_27 | 3 | -83.775 | 171.828 | *Lecidella* sp. nov2 | MK208756 | MK205051 | MK226994 | *Tr_S02* | MK226889 | MK226777 | MK227092 |
| MAF_MK1_31 | 3 | -83.775 | 171.828 | *Lecidea cancriformis* Dodge & Baker | MK208757 | MK205052 | MK226995 | *Tr_I01* | MK226890 | - | - |
| MAF_MK1_37 | 3 | -83.775 | 171.828 | *Lecidea cancriformis* Dodge & Baker | MK208758 | MK205053 | MK226996 | *Tr_I01* | MK226891 | - | - |
| MAF_MK1_38 | 3 | -83.775 | 171.828 | *Lecidea cancriformis* Dodge & Baker | MK208759 | MK205054 | - | *Tr_S02* | MK226892 | MK226778 | MK227093 |
| MAF_MK1_40 | 3 | -83.775 | 171.828 | *Lecanora fuscobrunnea* Dodge & Baker | MK208760 | MK205055 | MK226997 | *Tr_A02* | MK226893 | MK226754 | MK227094 |
| MAF_MK1_43 | 3 | -83.775 | 171.828 | *Lecidella* sp. nov2 | MK208761 | MK205056 | - | *Tr_S02* | MK226894 | MK226779 | MK227095 |
| MAF_MK1_48 | 3 | -83.775 | 171.828 | *Lecidea cancriformis* Dodge & Baker | MK208762 | MK205057 | MK226998 | *Tr_S02* | MK226895 | MK226780 | MK227096 |
| MAF_MK1_49 | 3 | -83.775 | 171.828 | *Lecidella* sp. nov2 | MK208763 | MK205058 | MK226999 | *Tr_S02* | MK226896 | MK226799 | MK227097 |
| MAF_MK1_55 | 3 | -83.775 | 171.828 | *Lecidea cancriformis* Dodge & Baker | MK208764 | MK205059 | MK227000 | - | - | - | - |
| MAF_MK1_56 | 3 | -83.775 | 171.828 | *Lecanora fuscobrunnea* Dodge & Baker | MK208765 | MK205060 | MK227001 | *Tr_S02* | MK226897 | MK226781 | MK227098 |
| MAF_MK1_63 | 3 | -83.775 | 171.828 | *Lecanora physciella* (Darb.) Hertel | MK208766 | MK205061 | MK227002 | *Tr_S02* | MK226898 | MK226789 | MK227099 |
| MAF_Sancho | 3 | -83.761 | 172.755 | *Lecidea cancriformis* Dodge & Baker | GU074439 | GU074489 | MK227003 | *Tr_A04a* | JN204838 | - | - |
| MAF_Sancho | 3 | -83.761 | 172.755 | *Carbonea* sp. 2 | MK208767 | - | - | *Tr_A04a* | JN204839 | - | - |
| T33335 | 3 | -83.803 | 172.207 | *Lecidella* sp. nov2 | MK208768 | MK205062 | - | *Tr_S18* | MK226899 | - | - |
| T33338 | 3 | -83.803 | 172.207 | *Lecidella* sp. nov2 | MK208769 | MK205063 | - | *Tr_S02* | MK226900 | - | - |
| T33346 | 3 | -83.803 | 172.207 | *Lecanora physciella* (Darb.) Hertel | JN873878 | - | - | - | - | - | - |
| T33348 | 3 | -83.803 | 172.207 | *Lecidea cancriformis* Dodge & Baker | - | MK205064 | - | - | - | - | - |
| Voucher ID | Area | Latitude | Longitude | Species name | Accession numbers |  |  | OTU ID | Accession numbers |  |  |
|  |  |  |  |  | nrITS | mtSSU | RPB1 |  | nrITS2 | psbJ-L | COX2 |
| T33446 | 3 | -83.828 | 172.749 | *Lecidea cancriiformis* Dodge & Baker | - | MK205065 | - | *Tr*_S18 | MK226901 | - | - |
| T33449 | 3 | -83.761 | 172.755 | *Lecidella siplei* Dodge & Baker | JN873897 | - | - | *Tr*_A02 | JN204729 | - | - |
| T33456 | 3 | -83.761 | 172.755 | *Lecidea andersonii* Filson | - | MK205066 | MK227004 | *Tr*_A02 | - | - | MK227100 |
| T33457 | 3 | -83.761 | 172.755 | *Lecidella siplei* Dodge & Baker | JN873898 | - | - | *Tr*_A02 | JN204731 | - | - |
| T35540 | 4a | -79.842 | 159.363 | *Lecidella siplei* Dodge & Baker | MK208770 | - | - | - | - | - | - |
| T35544 | 4a | -79.838 | 159.341 | *Lecanora fuscobrunnea* Dodge & Baker | JN873873 | - | - | - | - | - | - |
| T35559 | 4a | -79.838 | 159.221 | *Lecidea cancriiformis* Dodge & Baker | MK208771 | MK205067 | MK227005 | *Tr*_S18 | - | MK285375 | - |
| T35604 | 4a | -79.836 | 159.317 | *Lecidea cancriiformis* Dodge & Baker | EU257671 | GU074480 | MK227006 | *Tr*_S18 | JN204749 | - | - |
| T35620 | 4a | -79.835 | 159.385 | *Lecidea cancriiformis* Dodge & Baker | EU257672 | - | - | *Tr*_I01 | JN204750 | - | - |
| T35622 | 4a | -79.835 | 159.392 | *Lecidea cancriiformis* Dodge & Baker | MK208772 | MK205068 | MK227007 | - | - | - | - |
| T35647 | 4a | -79.851 | 159.341 | *Carbonea* sp. 2 | JN873866 | - | - | *Tr*_S02 | JN204751 | - | - |
| T35650 | 4a | -79.851 | 159.341 | *Carbonea* sp. 2 | JN873867 | - | MK227008 | *Tr*_S18 | JN204752 | - | - |
| T35662 | 4a | -79.840 | 159.332 | *Lecidea cancriiformis* Dodge & Baker | EU257673 | - | - | *Tr*_I01 | JN204753 | - | - |
| T35664 | 4a | -79.851 | 159.341 | *Lecanora fuscobrunnea* Dodge & Baker | JN873874 | - | - | - | - | - | - |
| T35686 | 4a | -79.842 | 159.363 | *Carbonea* sp. 2 | JN873868 | - | - | - | - | - | - |
| T42988 | 4b | -79.889 | 156.764 | *Lecidea cancriiformis* Dodge & Baker | GU074435 | GU074481 | MK227009 | *Tr*_S18 | MK226902 | MK226818 | MK227101 |
| T42990 | 4b | -79.926 | 156.890 | *Lecidea cancriiformis* Dodge & Baker | GU170841 | - | MK227010 | *Tr*_S18 | JN204770 | MK226819 | MK227102 |
| T42991 | 4b | -79.917 | 156.759 | *Lecidea cancriiformis* Dodge & Baker | GU170842 | MK205069 | MK227011 | *Tr*_S18 | MK226903 | - | - |
| T42992 | 4b | -79.917 | 156.751 | *Lecidea cancriiformis* Dodge & Baker | GU074436 | GU074482 | MK227012 | *Tr*_S18 | JN204771 | MK226820 | MK227103 |
| T42994 | 4b | -79.879 | 157.539 | *Lecanora fuscobrunnea* Dodge & Baker | GU170839 | MK205070 | MK227013 | *Tr*_S18 | MK226904 | - | - |
| T44625 | 4a | -79.868 | 159.360 | *Lecanora fuscobrunnea* Dodge & Baker | MK208773 | MK205071 | MK227014 | *Tr*_A02 | MK226905 | MK226755 | MK227104 |
| T44626 | 4a | -79.882 | 159.361 | *Carbonea* sp. URm1 | JN873865 | - | MK227015 | *Tr*_I01 | JN204797 | - | - |
| T44628 | 4a | -79.865 | 159.352 | *Lecanora fuscobrunnea* Dodge & Baker | JN873875 | MK205072 | MK227016 | *Tr*_S02 | - | MK226804 | MK227105 |
| T44632 | 4a | -79.868 | 159.352 | *Lecanora fuscobrunnea* Dodge & Baker | MK208774 | MK205073 | MK227017 | *Tr*_I01 | JN204800 | - | - |
| T44633 | 4a | -79.868 | 159.358 | *Lecidea cancriiformis* Dodge & Baker | MK208775 | MK205074 | - | *Tr*_S02 | - | MK226817 | MK227106 |
| T44634 | 4a | -79.869 | 159.341 | *Lecidea cancriiformis* Dodge & Baker | GU074434 | GU074486 | MK227018 | *Tr*_S18 | JN204801 | KF907601 | - |
| T44636 | 4a | -79.869 | 159.358 | *Lecidella greenii* Ruprecht & Türk | MK208776 | - | - | *Tr*_A02 | MK226906 | - | - |
| T44638 | 4a | -79.869 | 159.358 | *Lecidea cancriiformis* Dodge & Baker | - | MK205075 | - | *Tr*_S18 | MK226907 | MK226827 | MK227107 |
| T44640 | 4a | -79.869 | 159.358 | *Lecidea cancriiformis* Dodge & Baker | MK208777 | MK205076 | MK227019 | *Tr*_S02 | MK226908 | MK226809 | MK227108 |
| T44641 | 4a | -79.866 | 159.365 | *Carbonea* sp. 2 | JN873871 | - | MK227020 | *Tr*_A04a | JN204803 | KF907602 | - |
| T44643 | 4a | -79.859 | 159.238 | *Lecidea cancriiformis* Dodge & Baker | MK208778 | MK205077 | - | *Tr*_S18 | - | MK226831 | MK227109 |
| T44645 | 4a | -79.857 | 159.283 | *Lecidea cancriiformis* Dodge & Baker | MK208779 | MK205078 | MK227021 | *Tr*_S02 | MK226909 | MK226790 | MK227110 |
| T44646 | 4a | -79.857 | 159.283 | *Lecidea cancriiformis* Dodge & Baker | MK208780 | MK205079 | - | *Tr*_S02 | MK226910 | MK226793 | MK227111 |
| Voucher ID | Area | Latitude | Longitude | Species name | Accession numbers |  |  | OTU ID | Accession numbers |  |  |
|  |  |  |  |  | nrITS | mtSSU | RPB1 |  | nrITS2 | psbJ-L | COX2 |
| T44647 | 4a | -79.858 | 159.288 | *Lecidea cancriiformis* Dodge & Baker | MK208781 | MK205080 | - | Tr_S02 | MK226911 | MK226794 | MK227112 |
| T44648 | 4a | -79.858 | 159.288 | *Lecidea cancriiformis* Dodge & Baker | MK208782 | MK205081 | - | - | - | - | - |
| T44649 | 4a | -79.858 | 159.288 | *Lecidea cancriiformis* Dodge & Baker | MK208783 | MK205082 | - | Tr_S02 | MK226912 | MK226791 | MK227113 |
| T44650 | 4a | -79.858 | 159.288 | *Lecanora fuscobrunnea* Dodge & Baker | MK208784 | MK205083 | MK227022 | Tr_I01 | MK285376 | - | - |
| T44651 | 4a | -79.858 | 159.288 | *Lecidea cancriiformis* Dodge & Baker | MK208785 | MK205084 | - | Tr_S02 | MK226913 | MK226795 | MK227114 |
| T44652 | 4a | -79.858 | 159.288 | *Lecidea cancriiformis* Dodge & Baker | MK208786 | MK205085 | - | Tr_S02 | MK226914 | MK226796 | MK227115 |
| T44655 | 4a | -79.874 | 159.339 | *Carbonea* sp. 2 | MK208787 | - | MK227023 | Tr_S02 | MK226915 | MK226805 | MK227116 |
| T44656 | 4a | -79.863 | 159.373 | *Lecidea cancriiformis* Dodge & Baker | MK208788 | MK205086 | - | Tr_S02 | MK226916 | MK226797 | MK227117 |
| T44657 | 4a | -79.863 | 159.373 | *Lecanora fuscobrunnea* Dodge & Baker | JN873876 | MK205087 | MK227024 | Tr_S02 | JN204804 | MK226806 | MK227118 |
| T44659 | 4a | -79.864 | 159.368 | *Lecidea cancriiformis* Dodge & Baker | MK208789 | MK205088 | - | Tr_S02 | MK226917 | MK226798 | MK227119 |
| T44665 | 4a | -79.869 | 159.345 | *Lecidea cancriiformis* Dodge & Baker | MK208790 | MK205089 | - | Tr_S02 | MK226918 | MK226814 | MK227120 |
| T44666 | 4a | -79.869 | 159.345 | *Lecidea cancriiformis* Dodge & Baker | MK208791 | MK205090 | - | - | - | - | - |
| T44667 | 4a | -79.869 | 159.345 | *Lecidea cancriiformis* Dodge & Baker | MK208792 | MK205091 | MK227025 | Tr_S02 | MK226919 | MK226782 | MK227121 |
| T44669 | 4a | -79.863 | 159.378 | *Lecidea cancriiformis* Dodge & Baker | MK208793 | - | - | Tr_S02 | MK226920 | MK226768 | MK227122 |
| T44670 | 4a | -79.863 | 159.378 | *Lecidea cancriiformis* Dodge & Baker | MK208794 | MK205092 | - | Tr_I17 | MK226921 | - | - |
| T44674 | 4a | -79.877 | 159.326 | *Lecidea cancriiformis* Dodge & Baker | MK208795 | - | - | Tr_S02 | MK226923 | MK226783 | MK227124 |
| T44675 | 4a | -79.877 | 159.326 | *Lecanora fuscobrunnea* Dodge & Baker | - | MK205093 | - | - | JN204806 | - | - |
| T44676 | 4a | -79.883 | 159.348 | *Lecidea cancriiformis* Dodge & Baker | MK208796 | MK205094 | MK227026 | Tr_S18 | MK226924 | MK226832 | MK227125 |
| T44677 | 4a | -79.883 | 159.348 | *Lecidea cancriiformis* Dodge & Baker | MK208797 | MK205095 | MK227027 | Tr_A02 | MK226925 | MK226756 | MK227126 |
| T44679 | 4a | -79.877 | 159.331 | *Lecidea cancriiformis* Dodge & Baker | MK208798 | MK205096 | - | Tr_S02 | MK226927 | MK226769 | MK227128 |
| T44687 | 4a | -79.863 | 159.379 | *Lecanora fuscobrunnea* Dodge & Baker | MK208799 | - | - | - | - | - | MK227130 |
| T44688 | 4a | -79.863 | 159.382 | *Lecanora fuscobrunnea* Dodge & Baker | JN873877 | MK205097 | MK227028 | Tr_S02 | JN204807 | MK226807 | MK227131 |
| T44690 | 4a | -79.863 | 159.382 | *Lecidea cancriiformis* Dodge & Baker | MK208800 | MK205098 | - | Tr_S02 | MK226930 | MK226786 | MK227133 |
| T44692 | 4a | -79.866 | 159.366 | *Lecidea cancriiformis* Dodge & Baker | MK208801 | MK205099 | MK227029 | Tr_S02 | JN204809 | KF907603 | - |
| T44694 | 4b | -79.755 | 158.503 | *Lecidea cancriiformis* Dodge & Baker | MK208802 | MK205100 | - | Tr_I01 | MK226931 | - | - |
| T44695 | 4b | -79.761 | 158.503 | *Lecanora fuscobrunnea* Dodge & Baker | MK208803 | MK205101 | MK227030 | - | - | - | - |
| T44697 | 4b | -79.758 | 158.598 | *Lecanora fuscobrunnea* Dodge & Baker | MK208804 | MK205102 | - | Tr_S02 | MK226932 | MK226770 | MK227134 |
| T44698 | 4b | -79.755 | 158.627 | *Lecidea cancriiformis* Dodge & Baker | MK208805 | MK205103 | MK227031 | Tr_S18 | MK226933 | MK226825 | MK227135 |
| T44699 | 4b | -79.762 | 158.636 | *Lecanora fuscobrunnea* Dodge & Baker | MK208806 | MK205104 | MK227032 | Tr_A02 | MK226934 | MK226758 | MK227136 |
| T44700 | 4b | -79.762 | 158.637 | *Lecanora fuscobrunnea* Dodge & Baker | MK208807 | MK205105 | MK227033 | Tr_I01 | MK226935 | - | - |
| T44701 | 4b | -79.757 | 158.608 | *Lecidea cancriiformis* Dodge & Baker | MK208808 | MK205106 | - | Tr_S18 | MK226936 | MK226833 | MK227137 |
| T44702 | 4b | -79.756 | 158.610 | *Lecanora fuscobrunnea* Dodge & Baker | MK208809 | MK205107 | MK227034 | Tr_A02 | MK226937 | MK226759 | MK227138 |
| T44703 | 4b | -79.756 | 158.614 | *Lecidea cancriiformis* Dodge & Baker | - | MK205108 | MK227035 | Tr_S18 | MK226938 | MK226828 | MK227139 |
|  |  |  |  | Species name | Accession numbers |  |  | OTU ID | Accession numbers |  |  |
|  |  |  |  |  | nrITS | mtSSU | RPB1 |  | nrITS2 | psbJ-L | COX2 |
| T44704 | 4b | -79.756 | 158.617 | *Lecidea cancriiformis* Dodge & Baker | - | MK205109 | - | Tr_S18 | MK226939 | MK226829 | MK227140 |
| T44705 | 4b | -79.756 | 158.611 | *Lecanora fuscobrunnea* Dodge & Baker | MK208810 | - | - | Tr_S02 | MK226940 | MK226811 | MK227141 |
| T44707 | 4b | -79.758 | 158.598 | *Lecidea cancriiformis* Dodge & Baker | MK208811 | MK205110 | MK227036 | Tr_S18 | MK226941 | MK226834 | MK227142 |
| T44708 | 4b | -79.753 | 158.549 | *Lecidea cancriiformis* Dodge & Baker | MK208812 | MK205111 | - | Tr_S18 | - | MK226823 | MK227143 |
| T44709 | 4b | -79.753 | 158.541 | *Lecidea cancriiformis* Dodge & Baker | MK208813 | MK205112 | MK227037 | Tr_S18 | MK226942 | MK226826 | MK227144 |
| T44712 | 4b | -79.759 | 158.507 | *Lecidea cancriiformis* Dodge & Baker | GU074438 | GU074487 | MK227038 | Tr_S18 | JN204811 | - | - |
| T44713 | 4b | -79.759 | 158.511 | *Lecidea cancriiformis* Dodge & Baker | MK208814 | - | - | Tr_S18 | MK226945 | MK226836 | MK227147 |
| T44714 | 4b | -79.763 | 158.497 | *Lecidea cancriiformis* Dodge & Baker | MK208815 | MK205113 | - | Tr_S18 | MK226946 | MK226830 | MK227148 |
| T44715 | 4b | -79.758 | 158.602 | *Lecidea cancriiformis* Dodge & Baker | - | MK205114 | MK227039 | Tr_S18 | MK226947 | MK226837 | MK227149 |
| T44716 | 4b | -79.758 | 158.606 | *Lecidea cancriiformis* Dodge & Baker | - | MK205115 | MK227040 | Tr_S18 | MK226948 | MK226841 | MK227150 |
| T44717 | 4b | -79.755 | 158.620 | *Lecanora fuscobrunnea* Dodge & Baker | MK208816 | MK205116 | MK227041 | Tr_S02 | MK226949 | MK226787 | MK227151 |
| T44719 | 4b | -79.924 | 156.814 | *Lecanora fuscobrunnea* Dodge & Baker | MK208817 | MK205117 | MK227042 | Tr_S18 | MK226950 | MK226821 | MK227152 |
| T44720 | 4b | -79.949 | 156.789 | *Lecanora fuscobrunnea* Dodge & Baker | GU170840 | MK205118 | MK227043 | Tr_S02 | MK226951 | MK226808 | MK227153 |
| T44721 | 4b | -79.754 | 158.636 | *Lecidea cancriiformis* Dodge & Baker | MK208818 | MK205119 | MK227044 | Tr_S18 | MK226952 | MK226842 | MK227154 |
| T44723 | 4b | -79.878 | 157.527 | *Lecidea cancriiformis* Dodge & Baker | MK208819 | MK205120 | - | Tr_S18 | MK226953 | MK226838 | MK227155 |
| T44727 | 4b | -79.892 | 157.524 | *Lecidea cancriiformis* Dodge & Baker | GU074437 | GU074488 | - | Tr_S18 | JN204812 | - | - |
| T44787 | 4b | -79.929 | 156.705 | *Lecidea cancriiformis* Dodge & Baker | MK208820 | MK205122 | - | Tr_S18 | MK226955 | MK226839 | MK227157 |

**Supplementary Table S3.** Additional samples taken from Perez-Ortega et al. (2012) (Perez-Ortega et al. 2012) and used in this study, with information on collecting localities and Genbank accession numbers.

| Laboratory code | Area | Latitude | Longitude | Mycobiont |  | Associated green micro algae (*Trebouxia*) |  |
| --- | --- | --- | --- | --- | --- | --- | --- |
|  |  |  |  | Species name | Accession numbers nrITS | OTU ID | Accession numbers nrITS |
| s106 | 5 | -78.113 | 163.782 | *Lecanora* sp. 2 | JX036037 | Tr_A02 | JX036159 |
| s113 | 5 | -78.114 | 163.854 | *Lecidea cancriiformis* Dodge & Baker | JX036044 | Tr_S15 | JX036166 |
| s114 | 5 | -78.114 | 163.854 | *Lecidea cancriiformis* Dodge & Baker | JX036045 | Tr_A02 | JX036167 |
| s115 | 5 | -78.066 | 163.870 | *Lecidella greenii* Ruprecht & Türk | JX036046 | Tr_A02 | JX036168 |
| s120 | 5 | -78.083 | 163.768 | *Lecidea polypycnidophora* Ruprecht & Türk | JX036051 | Tr_A02 | JX036172 |
| s121 | 5 | -78.024 | 163.900 | *Carbonea vorticosa* (Flörke) Hertel | JX036052 | Tr_A02 | JX036173 |
| s122 | 5 | -78.110 | 163.787 | *Carbonea vorticosa* (Flörke) Hertel | JX036053 | Tr_A02 | JX036174 |
| s123 | 5 | -78.110 | 163.787 | *Carbonea vorticosa* (Flörke) Hertel | JX036054 | Tr_A02 | JX036175 |

|  |  |  |  | Mycobiont |  | Associated green micro algae (*Trebouxia*) |  |
| --- | --- | --- | --- | --- | --- | --- | --- |
| Laboratory code | Area | Latitude | Longitude | Species name | Accession numbers nrITS | OTU ID | Accession numbers nrITS |
| s124 | 5 | -78.057 | 163.844 | *Rhizoplaca macleanii* (Dodge) Castello | JX036055 | Tr_A02 | JX036176 |
| s125 | 5 | -78.057 | 163.844 | *Rhizoplaca macleanii* (Dodge) Castello | JX036056 | Tr_A02 | JX036177 |
| s171 | 5 | -78.036 | 163.837 | *Lecanora* sp. 2 | JX036076 | Tr_A02 | JX036197 |
| s173 | 5 | -78.114 | 163.854 | *Lecanora* sp. 2 | JX036078 | Tr_A02 | JX036199 |
| s175 | 5 | -78.033 | 163.849 | *Lecanora* sp. 2 | JX036080 | Tr_A02 | JX036201 |
| s179 | 5 | -78.063 | 163.809 | *Lecanora* sp. 2 | JX036084 | Tr_A02 | JX036205 |
| s181 | 5 | -78.111 | 163.858 | *Lecidella greenii* Ruprecht & Türk | JX036086 | Tr_A02 | JX036207 |
| s190 | 5 | -78.025 | 163.899 | *Lecanora* sp. 3 | JX036095 | Tr_A02 | JX036216 |
| s191 | 5 | -78.025 | 163.900 | *Lecanora* sp. 3 | JX036096 | Tr_A02 | JX036217 |
| s192 | 5 | -78.030 | 163.834 | *Lecidella greenii* Ruprecht & Türk | JX036097 | Tr_A02 | JX036218 |
| s197 | 5 | -78.061 | 163.791 | *Rhizoplaca macleanii* (Dodge) Castello | JX036101 | Tr_A02 | JX036222 |
| s198 | 5 | -78.061 | 163.791 | *Rhizoplaca macleanii* (Dodge) Castello | JX036102 | Tr_A02 | JX036223 |
| s201 | 5 | -78.024 | 163.900 | *Rhizoplaca macleanii* (Dodge) Castello | JX036105 | Tr_A02 | JX036226 |
| s202 | 5 | -78.027 | 163.839 | *Lecanora* cf. *mons-nivis* Darbishire | JX036106 | Tr_A02 | JX036227 |
| s203 | 5 | -78.027 | 163.839 | *Lecidella greenii* Ruprecht & Türk | JX036107 | Tr_A02 | JX036228 |
| s205 | 5 | -78.068 | 163.861 | *Rhizoplaca macleanii* (Dodge) Castello | JX036108 | Tr_A02 | JX036229 |
| s206 | 5 | -78.068 | 163.861 | *Rhizoplaca macleanii* (Dodge) Castello | JX036109 | Tr_A02 | JX036230 |
| s207 | 5 | -78.068 | 163.861 | *Rhizoplaca macleanii* (Dodge) Castello | JX036110 | Tr_A02 | JX036231 |
| s208 | 5 | -78.068 | 163.861 | *Rhizoplaca macleanii* (Dodge) Castello | JX036111 | Tr_A02 | JX036232 |
| s209 | 5 | -78.068 | 163.861 | *Rhizoplaca macleanii* (Dodge) Castello | JX036112 | Tr_A02 | JX036233 |
| s212 | 5 | -78.034 | 163.845 | *Rhizoplaca macleanii* (Dodge) Castello | JX036115 | Tr_A02 | JX036236 |
| s213 | 5 | -78.034 | 163.845 | *Rhizoplaca macleanii* (Dodge) Castello | JX036116 | Tr_A02 | JX036237 |
| s214 | 5 | -78.034 | 163.845 | *Rhizoplaca macleanii* (Dodge) Castello | JX036117 | Tr_A02 | JX036238 |
| s215 | 5 | -78.070 | 163.711 | *Lecanora fuscobrunnea* Dodge & Baker | JX036118 | Tr_A02 | JX036239 |
| s230 | 5 | -78.034 | 163.845 | *Lecidella greenii* Ruprecht & Türk | JX036133 | Tr_A02 | JX036252 |
| s232 | 5 | -78.034 | 163.845 | *Rhizoplaca macleanii* (Dodge) Castello | JX036135 | Tr_A02 | JX036254 |
| s233 | 5 | -78.034 | 163.845 | *Rhizoplaca macleanii* (Dodge) Castello | JX036136 | Tr_A02 | JX036255 |
| s235 | 5 | -78.113 | 163.778 | *Rhizoplaca macleanii* (Dodge) Castello | JX036138 | Tr_A02 | JX036257 |
| s236 | 5 | -78.113 | 163.778 | *Rhizoplaca macleanii* (Dodge) Castello | JX036139 | Tr_A02 | JX036258 |
| s237 | 5 | -78.113 | 163.778 | *Rhizoplaca macleanii* (Dodge) Castello | JX036140 | Tr_A02 | JX036259 |
| s266 | 5 | -78.075 | 163.791 | *Lecidella greenii* Ruprecht & Türk | JX036141 | Tr_A02 | JX036260 |
| s271 | 5 | -78.047 | 164.104 | *Rhizoplaca macleanii* (Dodge) Castello | JX036145 | Tr_A02 | JX036264 |
| s272 | 5 | -78.035 | 163.978 | *Rhizoplaca macleanii* (Dodge) Castello | JX036146 | Tr_A02 | JX036265 |
| s273 | 5 | -78.030 | 163.949 | *Rhizoplaca macleanii* (Dodge) Castello | JX036147 | Tr_A02 | JX036266 |
| s274 | 5 | -78.036 | 163.990 | *Rhizoplaca macleanii* (Dodge) Castello | JX036148 | Tr_A02 | JX036267 |
| Laboratory code | Area | Latitude | Longitude | Species name | Accession numbers nrITS | OTU ID | Accession numbers nrITS |
| s300 | 5 | -78.030 | 163.834 | *Lecidella greenii* Ruprecht & Türk | JX036150 | *Tr_A02* | JX036269 |
| s301 | 5 | -78.030 | 163.834 | *Lecidella greenii* Ruprecht & Türk | JX036151 | *Tr_A02* | JX036270 |
| s95 | 5 | -78.024 | 163.900 | *Rhizoplaca macleanii* (Dodge) Castello | JX036152 | *Tr_A02* | JX036271 |

**Supplementary Table S4.** Additional samples taken from Wagner et al. (2020) (Wagner et al. 2020) and used in this study, with information on collecting localities and Genbank accession numbers.

|  |  |  |  | Mycobiont |  |  |  | Associated green micro algae (*Trebouxia*) |  |  |  |
| --- | --- | --- | --- | --- | --- | --- | --- | --- | --- | --- | --- |
|  |  |  |  |  | Accession numbers |  |  |  | Accession numbers |  |  |
| Voucher ID | Area | Latitude | Longitude | Species name | nrITS | mtSSU | *RPB1* | OTU ID | nrITS | psbJ-L | COX2 |
| T46643 | 5 | -78.031 | 163.865 | *Rhizoplaca macleanii* (Dodge) Castello | MK970663 | - | - | *Tr_A02* | MK970698 | - | - |
| T46647b | 5 | -78.033 | 163.898 | *Rhizoplaca macleanii* (Dodge) Castello | MK970665 | MN023039 | MN023053 | *Tr_A02* | MK970698 | - | - |
| T46651 | 5 | -78.028 | 163.851 | *Carbonea vorticosa* (Flörke) Hertel | MK970656 | - | - | - | - | - | - |
| T46659 | 5 | -78.023 | 163.903 | *Lecidella greenii* Ruprecht & Türk | MK970671 | - | MN023055 | *Tr_A02* | MK970698 | - | - |
| T46672 | 5 | -78.024 | 163.898 | *Rhizoplaca macleanii* (Dodge) Castello | MK970663 | - | - | *Tr_A02* | MK970698 | - | - |
| T46673 | 5 | -78.027 | 163.851 | *Rhizoplaca macleanii* (Dodge) Castello | MK970663 | - | - | *Tr_A02* | MK970699 | - | - |
| T46676 | 5 | -78.028 | 163.851 | *Lecidella greenii* Ruprecht & Türk | MK970671 | - | - | *Tr_A02* | MK970699 | - | - |
| T46677 | 5 | -78.027 | 163.848 | *Lecidea cancriformis* Dodge & Baker | MK970681 | - | - | *Tr_A02* | MK970696 | - | - |
| T46678 | 5 | -78.028 | 163.850 | *Rhizoplaca macleanii* (Dodge) Castello | MK970669 | - | - | *Tr_A02* | MK970699 | - | - |
| T46679 | 5 | -78.036 | 163.971 | *Rhizoplaca macleanii* (Dodge) Castello | MK970664 | MN023039 | - | *Tr_A02* | MK970696 | - | - |
| T46680 | 5 | -78.036 | 163.971 | *Lecidea polypycnidophora* Ruprecht & Türk | MK970663 | MN023043 | MN023053 | *Tr_A02* | MK970699 | - | - |
| T46681 | 5 | -78.032 | 163.951 | *Rhizoplaca macleanii* (Dodge) Castello | MK970663 | - | - | *Tr_A02* | MK970698 | - | - |
| T46684 | 5 | -78.044 | 163.986 | *Lecidea cancriformis* Dodge & Baker | MK970677 | - | - | *Tr_S02* | MK970693 | - | - |
| T46685 | 5 | -78.044 | 163.986 | *Lecidea cancriformis* Dodge & Baker | MK970677 | - | MN023056 | *Tr_S02* | MK970693 | - | MN023030 |
| T46701 | 5 | -78.020 | 163.805 | *Rhizoplaca macleanii* (Dodge) Castello | MK970663 | MN023034 | - | *Tr_A02* | MK970698 | - | - |
| T46706 | 5 | -78.028 | 163.821 | *Lecidella greenii* Ruprecht & Türk | MK970671 | - | MN023054 | *Tr_A02* | MK970698 | - | - |
| T46710 | 5 | -78.073 | 163.717 | *Rhizoplaca macleanii* (Dodge) Castello | MK970666 | - | - | *Tr_A02* | MK970698 | - | - |
| T46713 | 5 | -78.028 | 163.843 | *Lecidea polypycnidophora* Ruprecht & Türk | MK970663 | MN023043 | MN023061 | *Tr_A02* | MK970698 | - | - |
| T46716 | 5 | -78.040 | 163.802 | *Lecidea polypycnidophora* Ruprecht & Türk | MK970663 | MN023043 | MN023061 | *Tr_A02* | MK970698 | - | - |
| T46717 | 5 | -78.040 | 163.806 | *Lecidea polypycnidophora* Ruprecht & Türk | MK970663 | - | - | *Tr_A02* | MK970698 | - | - |
| T46718 | 5 | -78.040 | 163.807 | *Carbonea vorticosa* (Flörke) Hertel | MK970656 | - | - | *Tr_A02* | MK970698 | - | - |
| T46719 | 5 | -78.038 | 163.804 | *Carbonea* sp. URm1 | MK970657 | - | - | *Tr_A02* | MK970698 | - | - |

|  |  |  |  | Mycobiont |  |  | Associated green micro algae (*Trebouxia*) |  |  |  |  |
| --- | --- | --- | --- | --- | --- | --- | --- | --- | --- | --- | --- |
| Voucher ID | Area | Latitude | Longitude | Species name | Accession numbers |  |  | OTU ID | Accession numbers |  |  |
|  |  |  |  |  | nrITS | mtSSU | RPB1 |  | nrITS | psbJ-L | COX2 |
| T48769 | 5 | -78.126 | 163.700 | *Lecidella greenii* Ruprecht & Türk | MK970671 | - | - | Tr_A02 | MK970699 | - | - |
| T48770 | 5 | -78.127 | 163.690 | *Lecanora* sp. 3 | MK970659 | - | - | Tr_A02 | MK970701 | - | - |
| T48773 | 5 | -78.127 | 163.674 | *Lecidea polypycnidophora* Ruprecht & Türk | MK970663 | MN023043 | - | Tr_A02 | MK970699 | MN023065 | - |
| T48774 | 5 | -78.135 | 163.626 | *Rhizoplaca macleanii* (Dodge) Castello | MK970663 | - | - | Tr_A02 | MK970698 | - | - |
| T48776 | 5 | -78.165 | 163.753 | *Lecidea cancriformis* Dodge & Baker | MK970679 | MN023046 | MN023057 | Tr_A02 | MK970702 | - | - |
| T48777 | 5 | -78.166 | 163.755 | *Lecidea cancriformis* Dodge & Baker | MK970679 | MN023046 | MN023058 | Tr_S15 | MK970692 | - | MN023031 |
| T48778a | 5 | -78.149 | 163.769 | *Rhizoplaca macleanii* (Dodge) Castello | MK970663 | - | - | Tr_A02 | MK970698 | - | - |
| T48779 | 5 | -78.164 | 163.755 | *Rhizoplaca macleanii* (Dodge) Castello | MK970664 | MN023039 | - | Tr_A02 | MK970696 | - | - |
| T48781 | 5 | -78.128 | 163.620 | *Lecidea* sp. 6 | MK620097 | - | - | Tr_A02 | MK970699 | - | - |
| T48782 | 5 | -78.123 | 163.642 | *Lecidea cancriformis* Dodge & Baker | MK970679 | MN023046 | - | Tr_S18 | MK970695 | MN023070 | MN023032 |
| T48784 | 5 | -78.121 | 163.683 | *Lecidea polypycnidophora* Ruprecht & Türk | MK970663 | MN023043 | - | Tr_A02 | MK970699 | MN023065 | - |
| T48785 | 5 | -78.133 | 163.666 | *Lecidella greenii* Ruprecht & Türk | MK970671 | - | MN023054 | Tr_S15 | MK970692 | - | - |
| T48787 | 5 | -78.127 | 163.678 | *Lecidella greenii* Ruprecht & Türk | MK970671 | - | MN023054 | Tr_A02 | MK970699 | - | - |
| T48788 | 5 | -78.120 | 163.684 | *Carbonea vorticosa* (Flörke) Hertel | MK970656 | MN023033 | MN023050 | Tr_A02 | MK970699 | - | - |
| T48789 | 5 | -78.152 | 163.739 | *Carbonea* sp. 2 | MK970654 | - | MN023051 | Tr_S15 | MK970692 | - | MN023031 |
| T48790a | 5 | -78.120 | 163.682 | *Carbonea vorticosa* (Flörke) Hertel | MK970656 | MN023033 | - | Tr_A02 | MK970699 | - | - |
| T48790b | 5 | -78.120 | 163.682 | *Lecanora* sp. 3 | MK970659 | - | - | Tr_A02 | MK970703 | - | - |
| T48791a | 5 | -78.120 | 163.686 | *Lecidella greenii* Ruprecht & Türk | MK970671 | - | - | Tr_A02 | MK970699 | - | - |
| T48793a | 5 | -78.153 | 163.731 | *Lecidea cancriformis* Dodge & Baker | MK970677 | - | MN023056 | Tr_S15 | MK970692 | - | MN023031 |
| T48793b | 5 | -78.153 | 163.731 | *Carbonea* sp. 2 | MK970654 | - | - | Tr_S15 | MK970692 | - | MN023031 |
| T48794b | 5 | -78.151 | 163.735 | *Rhizoplaca macleanii* (Dodge) Castello | MK970663 | - | - | Tr_A02 | MK970698 | - | - |
| T48795a | 5 | -78.161 | 163.714 | *Rhizoplaca macleanii* (Dodge) Castello | MK970668 | - | - | Tr_A02 | MK970698 | - | - |
| T48797 | 5 | -78.156 | 163.689 | *Rhizoplaca macleanii* (Dodge) Castello | MK970667 | - | - | Tr_A02 | MK970698 | - | - |
| T48798 | 5 | -78.150 | 163.736 | *Rhizoplaca macleanii* (Dodge) Castello | MK970667 | - | - | Tr_A02 | MK970698 | - | - |
| T48799b | 5 | -78.145 | 163.620 | *Lecidea cancriformis* Dodge & Baker | MK970679 | - | - | Tr_A02 | MK970698 | - | - |
| T48799c | 5 | -78.145 | 163.620 | *Lecidea cancriformis* Dodge & Baker | MK970677 | - | - | Tr_S15 | MK970692 | - | - |
| T48800 | 5 | -78.146 | 163.631 | *Lecanora* sp. 2 | MK970662 | MN023036 | MN023052 | Tr_A02 | MK970698 | - | - |
| T48801a | 5 | -78.144 | 163.626 | *Lecanora* sp. 3 | MK970659 | - | - | Tr_A02 | MK970699 | - | - |
| T48801c | 5 | -78.144 | 163.626 | *Lecidella greenii* Ruprecht & Türk | MK970671 | MN023040 | - | Tr_A02 | MK970699 | - | - |
| T48803b | 5 | -78.148 | 163.630 | *Carbonea* sp. URm1 | MK970657 | MN023034 | - | Tr_A02 | MK970699 | - | - |
| T48804 | 5 | -78.148 | 163.630 | *Rhizoplaca macleanii* (Dodge) Castello | MK970670 | - | MN023053 | Tr_A02 | MK970699 | - | - |
| T48805 | 5 | -78.144 | 163.626 | *Lecidella greenii* Ruprecht & Türk | MK970671 | - | - | Tr_A02 | MK970698 | - | - |
| T48806 | 5 | -78.142 | 163.628 | *Carbonea* sp. URm1 | MK970657 | MN023034 | - | Tr_A02 | MK970699 | - | - |
| Voucher ID | Area | Latitude | Longitude | Species name | Accession numbers |  |  | OTU ID | Accession numbers |  |  |
|  |  |  |  |  | nrITS | mtSSU | RPB1 |  | nrITS | psbJ-L | COX2 |
| T48807 | 5 | -78.141 | 163.631 | *Lecidea andersonii* Filson | MK970673 | MN023042 | MN023060 | Tr_A02 | MK970702 | - | - |
| T48809 | 5 | -78.148 | 163.657 | *Lecidella greenii* Ruprecht & Türk | MK970671 | - | - | Tr_A02 | MK970699 | - | - |
| T48811a | 5 | -78.142 | 163.657 | *Lecidella greenii* Ruprecht & Türk | MK970671 | - | MN023055 | Tr_A02 | MK970698 | - | - |
| T48812a | 5 | -78.138 | 163.619 | *Lecidella greenii* Ruprecht & Türk | MK970671 | - | - | Tr_A02 | MK970699 | - | - |
| T48812b | 5 | -78.138 | 163.619 | *Lecidella greenii* Ruprecht & Türk | MK970671 | - | MN023055 | Tr_A02 | MK970699 | - | - |
| T48813 | 5 | -78.142 | 163.657 | *Lecidea* UCR1 | MK970675 | MN023044 | - | Tr_A02 | MK970699 | - | - |
| T48817a | 5 | -78.113 | 163.785 | *Rhizoplaca macleanii* (Dodge) Castello | MK970663 | - | - | Tr_A02 | MK970698 | - | - |
| T48817b | 5 | -78.113 | 163.785 | *Carbonea* sp. 2 | MK970655 | - | - | Tr_A02 | MK970702 | - | - |
| T48820 | 5 | -78.114 | 163.780 | *Carbonea* sp. 2 | MK970654 | - | - | Tr_A02 | MK970698 | - | - |
| T48821 | 5 | -78.114 | 163.779 | *Rhizoplaca macleanii* (Dodge) Castello | MK970665 | - | - | Tr_A02 | MK970698 | - | - |
| T48823a | 5 | -78.098 | 163.710 | *Carbonea vorticosa* (Flörke) Hertel | MK970656 | - | - | - | - | - | - |
| T48825 | 5 | -78.097 | 163.691 | *Carbonea* sp. 2 | MK970654 | MN023035 | - | Tr_A02 | MK970699 | - | - |
| T48826 | 5 | -78.097 | 163.717 | *Lecanora fuscobrunnea* Dodge & Baker | MK970661 | - | - | Tr_S02 | MK970694 | MN023068 | - |
| T48828a | 5 | -78.110 | 163.858 | *Lecidella greenii* Ruprecht & Türk | MK970671 | - | MN023054 | - | - | - | - |
| T48828b | 5 | -78.110 | 163.858 | *Lecidella greenii* Ruprecht & Türk | MK970671 | - | - | Tr_A02 | MK970698 | - | - |
| T48829 | 5 | -78.111 | 163.858 | *Lecidella greenii* Ruprecht & Türk | MK970671 | - | - | Tr_A02 | MK970699 | - | - |
| T48831 | 5 | -78.111 | 163.858 | *Carbonea vorticosa* (Flörke) Hertel | MK970656 | MN023033 | - | Tr_A02 | MK970699 | - | - |
| T48832 | 5 | -78.112 | 163.824 | *Carbonea* sp. 2 | MK970654 | - | MN023051 | Tr_A02 | MK970702 | - | - |
| T48836 | 5 | -78.099 | 163.778 | *Lecanora cf. mons-nivis* Darbishire | MK970658 | - | - | Tr_A02 | MK970699 | - | - |
| T48837 | 5 | -78.098 | 163.777 | *Lecidella greenii* Ruprecht & Türk | MK970671 | - | MN023055 | Tr_A02 | MK970699 | - | - |
| T48839 | 5 | -78.114 | 163.854 | *Lecidella greenii* Ruprecht & Türk | MK970671 | - | - | Tr_A02 | MK970699 | - | - |
| T48841a | 5 | -78.068 | 163.861 | *Rhizoplaca macleanii* (Dodge) Castello | MK970663 | - | - | Tr_A02 | MK970698 | - | - |
| T48841b | 5 | -78.068 | 163.861 | *Carbonea vorticosa* (Flörke) Hertel | MK970656 | - | MN023050 | Tr_A02 | MK970698 | - | - |
| T48843a | 5 | -78.066 | 163.870 | *Lecidea cancriformis* Dodge & Baker | MK970677 | - | - | Tr_S02 | MK970693 | - | - |
| T48843c | 5 | -78.066 | 163.870 | *Lecidea cancriformis* Dodge & Baker | MK970678 | - | - | - | - | - | - |
| T48843d | 5 | -78.066 | 163.870 | *Rhizoplaca macleanii* (Dodge) Castello | MK970663 | - | - | Tr_A02 | MK970699 | - | - |
| T48843e | 5 | -78.066 | 163.870 | *Rhizoplaca macleanii* (Dodge) Castello | MK970663 | - | - | Tr_A02 | MK970702 | - | - |
| T48844 | 5 | -78.067 | 163.863 | *Lecidella greenii* Ruprecht & Türk | MK970671 | - | - | Tr_A02 | MK970698 | - | - |
| T48851 | 5 | -78.074 | 163.793 | *Lecanora fuscobrunnea* Dodge & Baker | MK970660 | MN023037 | - | Tr_A02 | MK970698 | - | - |
| T48855b | 5 | -78.036 | 163.837 | *Lecidea cancriformis* Dodge & Baker | MK970680 | MN023046 | MN023059 | Tr_S15 | MK970692 | - | MN023031 |
| T48857a | 5 | -78.036 | 163.827 | *Lecidea* sp. 6 | MK970684 | MN023045 | - | Tr_A02 | MK970699 | - | - |
| T48858 | 5 | -78.025 | 163.975 | *Lecidella greenii* Ruprecht & Türk | MK970671 | - | - | - | - | - | - |
| T48859 | 5 | -78.025 | 163.986 | *Lecidea polypycnidophora* Ruprecht & Türk | MK970663 | - | MN023061 | Tr_A02 | MK970698 | - | - |
| Voucher ID | Area | Latitude | Longitude | Species name | Accession numbers |  |  | OTU ID | Accession numbers |  |  |
|  |  |  |  |  | nrITS | mtSSU | RPB1 |  | nrITS | psbJ-L | COX2 |
| T48860b | 5 | -78.036 | 163.836 | *Lecidea* sp. 6 | MK970684 | MN023045 | MN023064 | *Tr_A02* | MK970699 | - | - |
| T48861 | 5 | -78.024 | 163.892 | *Lecanora* sp. 3 | MK970659 | - | - | *Tr_A02* | MK970699 | - | - |
| T48862 | 5 | -78.037 | 163.978 | *Lecidea cancriformis* Dodge & Baker | MK970682 | - | - | - | - | - | - |
| T48864 | 5 | -78.039 | 163.989 | *Lecidea cancriformis* Dodge & Baker | MK970679 | - | - | *Tr_A02* | MK970702 | - | - |
| T48865a | 5 | -78.036 | 163.990 | *Rhizoplaca macleanii* (Dodge) Castello | MK970666 | - | - | *Tr_A02* | MK970698 | MN023067 | - |
| T48867a | 5 | -78.043 | 164.104 | *Rhizoplaca macleanii* (Dodge) Castello | MK970663 | - | - | *Tr_A02* | MK970696 | - | - |
| T48867c | 5 | -78.043 | 164.104 | *Rhizoplaca macleanii* (Dodge) Castello | MK970666 | - | - | *Tr_A02* | MK970702 | - | - |
| T48869 | 5 | -78.037 | 163.978 | *Lecidella greenii* Ruprecht & Türk | MK970671 | - | - | - | - | - | - |
| T48872 | 5 | -78.030 | 163.951 | *Lecidella greenii* Ruprecht & Türk | MK970671 | - | - | - | - | - | - |
| T48873 | 5 | -78.024 | 163.893 | *Lecidella greenii* Ruprecht & Türk | MK970671 | - | MN023054 | *Tr_A02* | MK970698 | - | - |
| T48874 | 5 | -78.024 | 163.899 | *Lecidea* UCR1 | MK970676 | MN023044 | MN023062 | *Tr_A02* | MK970697 | MN023066 | - |
| T48875 | 5 | -78.024 | 163.900 | *Lecidea lapicida* (Ach.) Ach. subsp. | MK970683 | - | - | *Tr_A02* | MK970699 | - | - |
| T48876a | 5 | -78.024 | 163.900 | *Lecidella greenii* Ruprecht & Türk | MK970671 | MN023040 | - | *Tr_A02* | MK970698 | - | - |
| T48876b | 5 | -78.024 | 163.900 | *Lecidella greenii* Ruprecht & Türk | MK970671 | - | MN023055 | *Tr_A02* | MK970699 | - | - |
| T48877 | 5 | -78.024 | 163.900 | *Carbonea vorticosa* (Flörke) Hertel | MK970656 | - | MN023050 | *Tr_A02* | MK970698 | - | - |
| T48879 | 5 | -78.057 | 163.747 | *Lecidella greenii* Ruprecht & Türk | MK970671 | - | MN023055 | - | - | - | - |
| T48880 | 5 | -78.058 | 163.740 | *Lecidea polypycnidophora* Ruprecht & Türk | MK970674 | - | - | - | - | - | - |
| T48881b | 5 | -78.061 | 163.791 | *Rhizoplaca macleanii* (Dodge) Castello | MK970666 | - | - | *Tr_A02* | MK970699 | - | - |
| T48882 | 5 | -78.057 | 163.817 | *Lecidea polypycnidophora* Ruprecht & Türk | MK970663 | - | - | *Tr_A02* | MK970699 | MN023065 | - |
| T48883b | 5 | -78.058 | 163.847 | *Lecidea* sp. 5 | MK620099 | - | MN023063 | *Tr_A02* | MK970700 | - | - |
| T48885 | 5 | -78.057 | 163.844 | *Rhizoplaca macleanii* (Dodge) Castello | MK970666 | - | - | *Tr_A02* | MK970698 | - | - |
| T48887a | 5 | -78.114 | 163.854 | *Lecidea cancriformis* Dodge & Baker | MK970677 | - | - | *Tr_S15* | MK970692 | - | - |
| T48887a | 5 | -78.070 | 163.711 | *Lecidea cancriformis* Dodge & Baker | MK970679 | MN023046 | - | *Tr_S15* | MK970692 | - | - |
| T48888b | 5 | -78.073 | 163.717 | *Lecanora* cf. *mons-nivis* Darbishire | MK970658 | - | - | *Tr_A02* | MK970699 | - | - |
| T48900 | 5 | -78.034 | 163.845 | *Rhizoplaca macleanii* (Dodge) Castello | MK970663 | - | - | *Tr_A02* | MK970698 | - | - |

**Supplementary Table S5.** Diversity metrics compared in this study, citations, descriptions and interpretation of each, and the used R functions.

| Metric | Definition | Citations | Description and interpretation of values | R functions, R package |
| --- | --- | --- | --- | --- |
| ***NRI*** | Net relatedness index | Webb, 2000; Webb, 2002 (Webb 2000; Webb et al. 2002) | Comparison of phylogenetic distances among all members of a community (pos. values = phylogenetic clustering; neg. values = phylogenetic evenness) | ses.mpd(), picante (Kembel et al. 2010) |
| ***PSR*** | Phylogenetic species richness | Helmus, 2007 (Helmus et al. 2007) | PSV (phylogenetic species variability; degree to which species in a community are phylogenetically related) multiplied by species richness SR (number of species in a sample); SR after discounting species relatedness (values range from 0 = increased relatedness to SR = decreased relatedness) | psd(), picante (Kembel et al. 2010) |
| ***J'*** | Pielou evenness index | Pielou, 1969 (Pielou 1969) | Measure of how evenly distributed abundance is numerically among the species that exist in a community (values range from 0 = no evenness to 1 = complete evenness) | diversity(), vegan (Oksanen et al. 2019) |
| ***1 - J'*** | 1 - Pielou evenness index |  | Values range from 0 = complete evenness to 1 = no evenness |  |

**Supplementary Table S6.** Diversity indices (left), specificity indices (middle) and BIO10, BIO12, elevation and latitude means (right) for the different mycobiont species and photobiont OTUs: *N*, number of sequences; *h*, number of haplotypes; *h / N*, ratio of *h* and *N*; *Hd*, haplotype diversity; *π*, nucleotide diversity; *NRI*, net relatedness index; *PSR*, phylogenetic species richness; *J’*, Pielou evenness index. (Note: the specificity indices were calculated for the respective symbiosis partners: *1 – J’* on the basis of species/OTUs, *NRI* and PSR on the basis of haplotypes. As a consequence, only samples where both mycobiont as well as photobiont could be identified are included.)

| Mycobiont species | *N* | *h* | *h* / *N* | *Hd* | $\pi$ | *NRI* | *PSR* | $1 - J'$ | BIO10 mean | BIO12 mean | Elevation mean (m. a. s. l.) | Latitude mean |
| --- | --- | --- | --- | --- | --- | --- | --- | --- | --- | --- | --- | --- |
| *Carbonea* sp. 2 | 13 | 3 | 0.231 | 0.564 | 0.0014 | 1.238 | 6.473 | 0.220 | -7.33 | 120.85 | 518.83 | -79.79 |
| *Carbonea* sp. URm1 | 5 | 2 | 0.400 | 0.400 | 0.0128 | -0.297 | 2.913 | 0.743 | -6.78 | 138.00 | 524.40 | -79.95 |
| *Carbonea vorticosa* | 11 | 1 | 0.091 | - | - | 1.471 | 2.000 | 1.000 | -6.27 | 143.73 | 436.09 | -78.08 |
| *Lecanora* cf. *mons-nivis* | 3 | 2 | 0.667 | 0.667 | 0.0000 | 1.413 | 2.000 | 1.000 | -6.30 | 137.33 | 389.33 | -78.07 |
| *Lecanora fuscobrunnea* | 31 | 6 | 0.194 | 0.546 | 0.0013 | -0.423 | 8.332 | 0.399 | -8.35 | 107.41 | 574.81 | -81.11 |
| *Lecanora physciella* | 3 | 2 | 0.667 | 0.667 | 0.0014 | 1.473 | 0.011 | 1.000 | -7.70 | 103.67 | 610.33 | -84.04 |
| *Lecanora* sp. 2 | 6 | 1 | 0.167 | - | - | 2.305 | 3.000 | 1.000 | -6.23 | 141.00 | 606.83 | -78.08 |
| *Lecanora* sp. 3 | 7 | 1 | 0.143 | - | - | 3.002 | 2.727 | 1.000 | -6.47 | 153.86 | 450.43 | -79.14 |
| *Lecidea andersonii* | 8 | 1 | 0.125 | - | - | -0.307 | 1.764 | 0.806 | -7.19 | 117.88 | 282.50 | -83.65 |
| *Lecidea cancriformis* | 93 | 18 | 0.194 | 0.797 | 0.0032 | -1.119 | 10.994 | 0.244 | -8.18 | 112.87 | 601.86 | -80.82 |
| *Lecidea lapicida* | 1 | 1 | 1.000 | - | - | - | - | 1.000 | -5.90 | 151.00 | 375.00 | -78.02 |
| *Lecidea polypycnidophora* | 10 | 1 | 0.100 | - | - | 1.471 | 2.000 | 1.000 | -6.22 | 141.10 | 411.20 | -78.06 |
| *Lecidea* sp. 5 | 1 | 1 | 1.000 | - | - | - | - | 1.000 | -6.50 | 172.00 | 671.00 | -78.06 |
| *Lecidea* sp. 6 | 3 | 2 | 0.667 | 0.667 | 0.0106 | - | - | 1.000 | -6.33 | 148.67 | 709.00 | -78.07 |
| *Lecidea* UCR1 | 2 | 1 | 0.500 | - | - | 1.450 | 2.000 | 1.000 | -6.25 | 140.00 | 466.00 | -78.08 |
| *Lecidella greenii* | 37 | 3 | 0.081 | 0.324 | 0.0007 | 1.933 | 4.600 | 0.929 | -6.33 | 144.95 | 488.57 | -78.53 |
| *Lecidella siplei* | 10 | 4 | 0.400 | 0.644 | 0.0012 | 2.974 | 2.916 | 1.000 | -7.52 | 136.80 | 322.30 | -84.21 |
| *Lecidella* sp. nov2 | 9 | 3 | 0.333 | 0.556 | 0.0073 | 1.643 | 0.403 | 0.821 | -7.90 | 100.33 | 787.67 | -83.87 |
| *Rhizoplaca macleanii* | 51 | 8 | 0.157 | 0.707 | 0.0019 | 4.209 | 4.860 | 1.000 | -6.18 | 149.92 | 684.53 | -78.07 |
| Photobiont OTU | *N* | *h* | *h* / *N* | *Hd* | $\pi$ | *NRI* | *PSR* | $1 - J'$ | BIO10 mean | BIO12 mean | Elevation mean (m. a. s. l.) | Latitude mean |
| *Tr_A02* | 165 | 14 | 0.085 | 0.702 | 0.0025 | 1.739 | 16.265 | 0.220 | -6.47 | 144.24 | 538.61 | -79.20 |
| *Tr_A04a* | 3 | 3 | 1.000 | 1.000 | 0.0023 | -0.759 | 3.000 | 0.784 | -8.70 | 116.67 | 491.00 | -82.46 |
| *Tr_I01* | 10 | 6 | 0.600 | 0.844 | 0.0084 | 0.036 | 1.955 | 0.695 | -8.12 | 98.60 | 555.70 | -81.19 |
| *Tr_I17* | 2 | 1 | 0.500 | - | - | - | - | 1.000 | -7.60 | 101.50 | 358.50 | -82.21 |
| *Tr_S02* | 58 | 8 | 0.138 | 0.573 | 0.0049 | 0.538 | 7.970 | 0.580 | -7.89 | 120.25 | 565.10 | -82.12 |
| *Tr_S15* | 10 | 1 | 0.100 | - | - | 0.578 | 2.596 | 0.728 | -6.16 | 139.10 | 689.10 | -78.12 |
| *Tr_S18* | 32 | 1 | 0.031 | - | - | 2.529 | 3.431 | 0.828 | -9.91 | 77.88 | 696.25 | -80.02 |

**Supplementary Table S7.** Network matrix giving the number of associations between the mycobiont species and photobiont OTUs.

| Mycobiont species | Photobiont OTU |  |  |  |  |  |  |
| --- | --- | --- | --- | --- | --- | --- | --- |
|  | Tr_A02 | Tr_A04a | Tr_I01 | Tr_I17 | Tr_S02 | Tr_S15 | Tr_S18 |
| *Carbonea* sp. 2 | 4 | 2 | - | - | 3 | 2 | 1 |
| *Carbonea* sp. URM1 | 4 | - | 1 | - | - | - | - |
| *Carbonea vorticosa* | 9 | - | - | - | - | - | - |
| *Lecanora* cf. *mons-nivis* | 3 | - | - | - | - | - | - |
| *Lecanora fuscobrunnea* | 11 | - | 3 | - | 11 | - | 2 |
| *Lecanora physciella* | - | - | - | - | 2 | - | - |
| *Lecanora* sp. 2 | 6 | - | - | - | - | - | - |
| *Lecanora* sp. 3 | 7 | - | - | - | - | - | - |
| *Lecidea andersonii* | 7 | - | - | - | 1 | - | - |
| *Lecidea cancriformis* | 7 | 1 | 6 | 2 | 34 | 7 | 28 |
| *Lecidea lapicida* | 1 | - | - | - | - | - | - |
| *Lecidea polypycnidophora* | 9 | - | - | - | - | - | - |
| *Lecidea* sp. 5 | 1 | - | - | - | - | - | - |
| *Lecidea* sp. 6 | 3 | - | - | - | - | - | - |
| *Lecidea* UCR1 | 2 | - | - | - | - | - | - |
| *Lecidella greenii* | 31 | - | - | - | - | 1 | - |
| *Lecidella siplei* | 9 | - | - | - | - | - | - |
| *Lecidella* sp. nov2 | - | - | - | - | 8 | - | 1 |
| *Rhizoplaca macleanii* | 51 | - | - | - | - | - | - |

## Supplementary Material 2: Figures

**Supplementary Figure S1.**
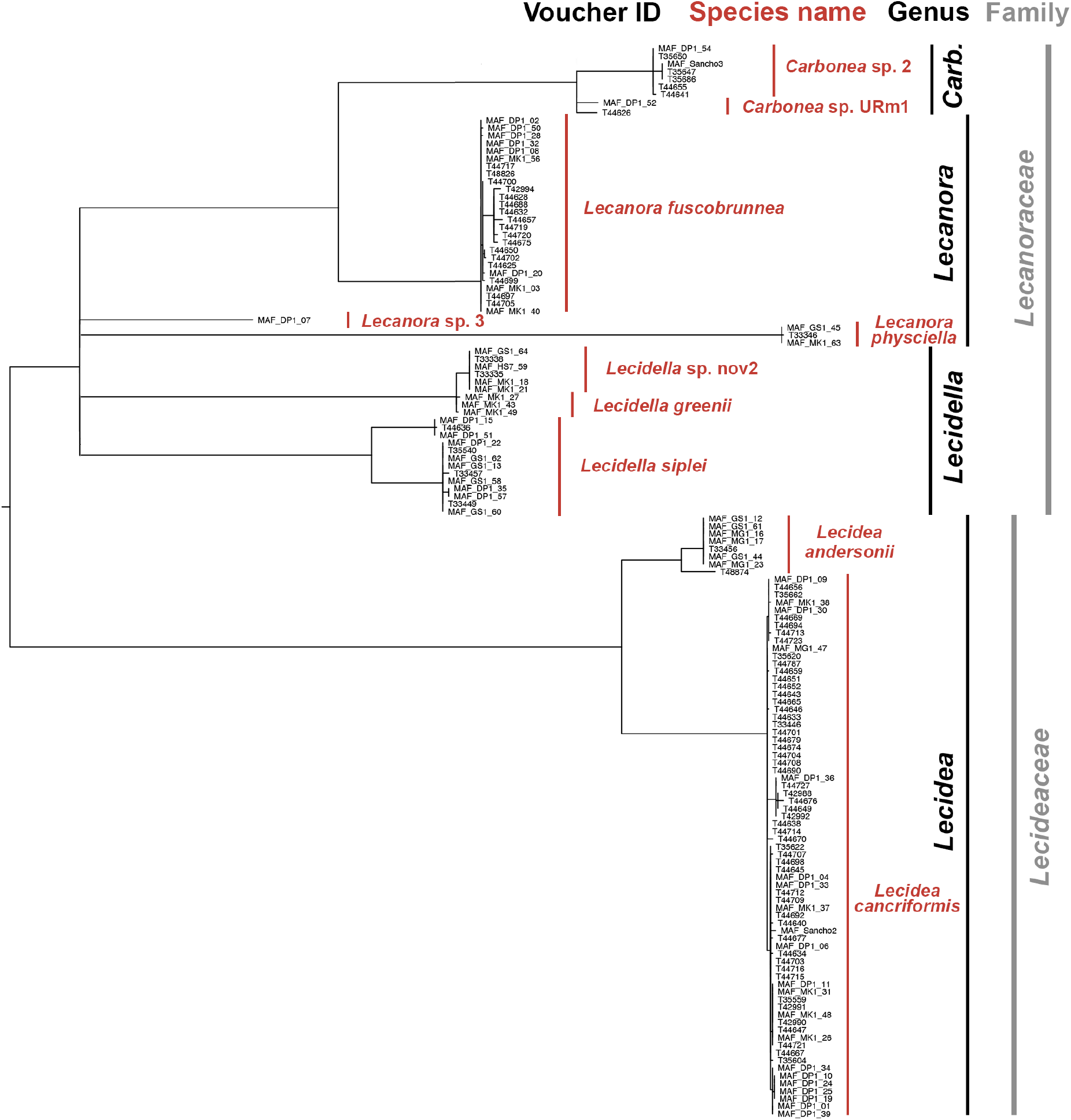
Phylogeny of mycobiont specimen based on multi-locus sequence data (nrITS, mtSSU and RPB1; calculated with IQ-TREE (Nguyen et al. 2014); branches with SH-aLRT < 80 % and UFboot < 95 % were collapsed).

**Supplementary Figure S2.**
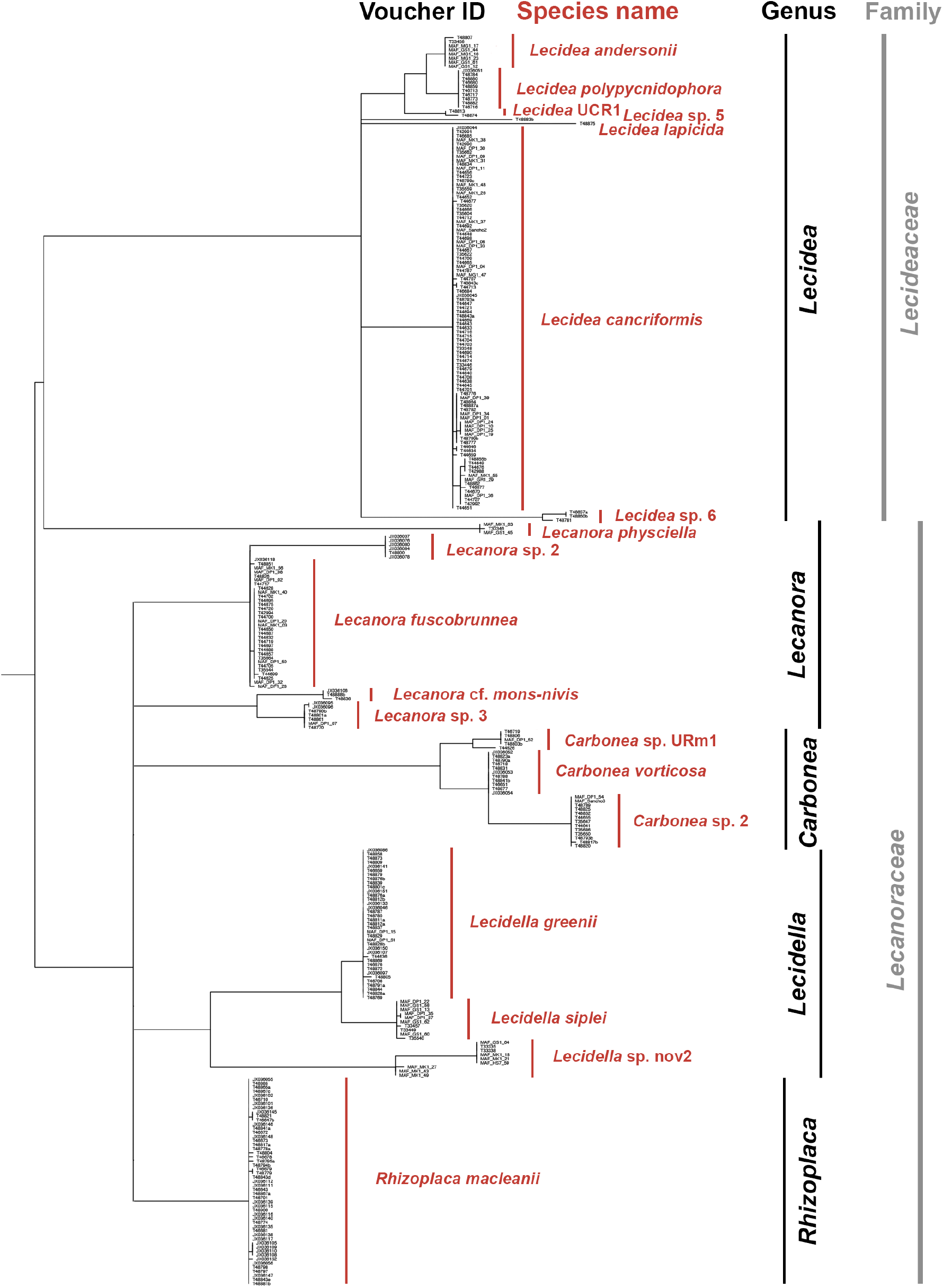
Phylogeny of all mycobiont specimen based on the marker nrITS (calculated with IQ-TREE (Nguyen et al. 2014); branches with SH-aLRT < 80 % and UFboot < 95 % were collapsed).

**Supplementary Figure S3.**
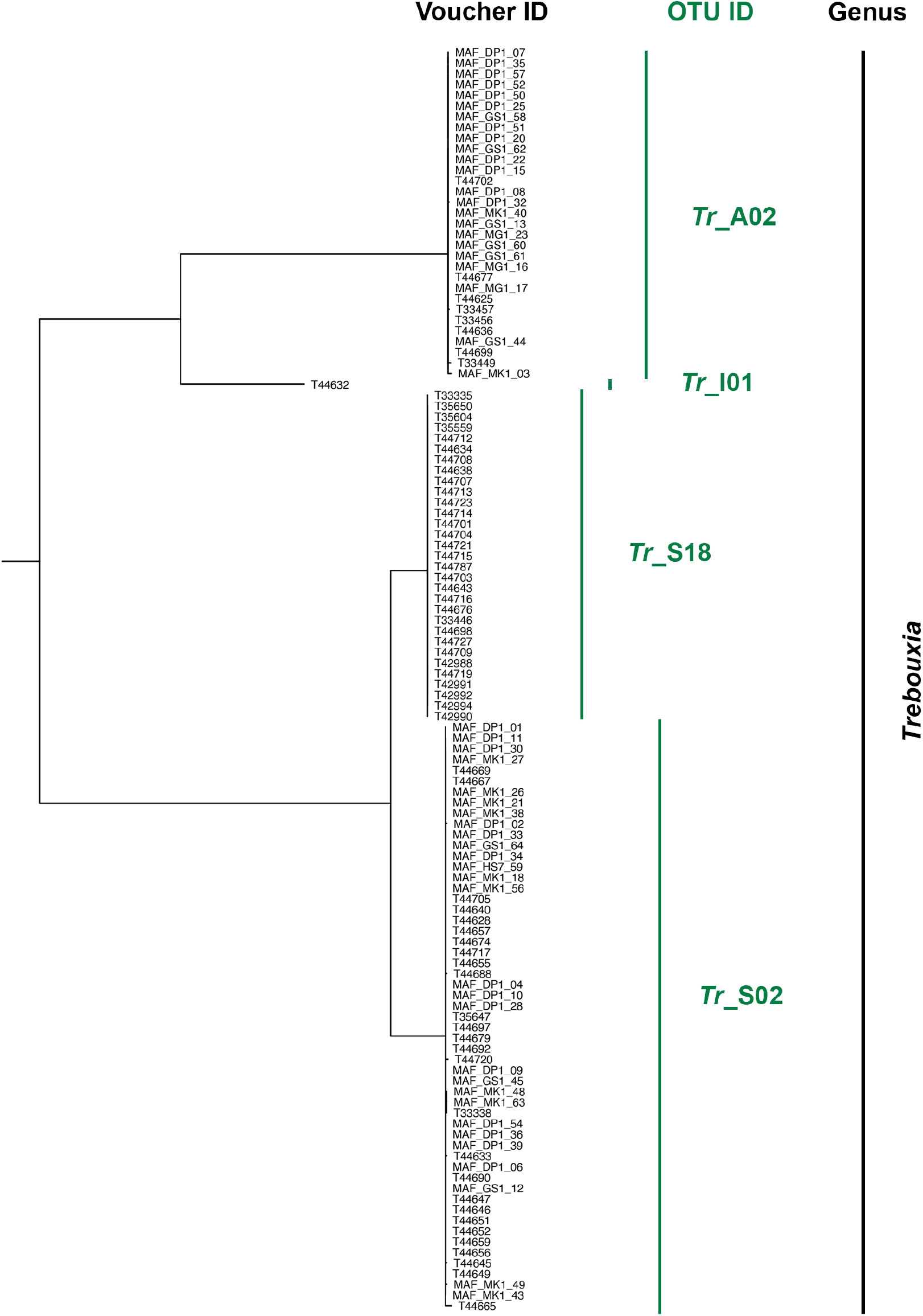
Phylogeny of photobiont specimen based on multi-locus sequence data (nrITS, psbJ-L and COX2; calculated with IQ-TREE (Nguyen et al. 2014); branches with SH-aLRT < 80 % and UFboot < 95 % were collapsed).

**Supplementary Figure S4.**
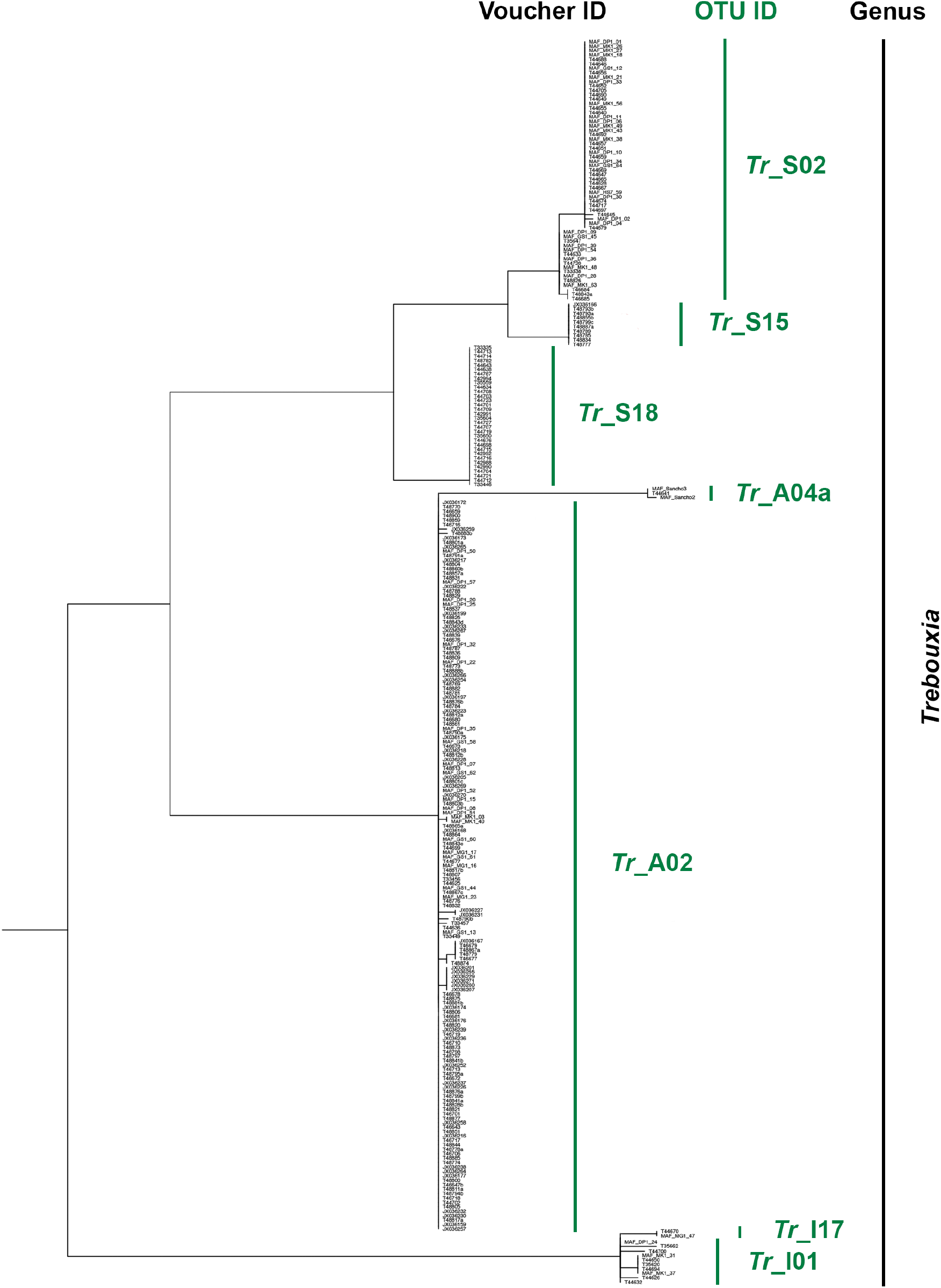
Phylogeny of all photobiont specimen based on the marker nrITS (calculated with IQ-TREE (Nguyen et al. 2014); branches with SH-aLRT < 80 % and UFboot < 95 % were collapsed).

**Supplementary Figure S5.**
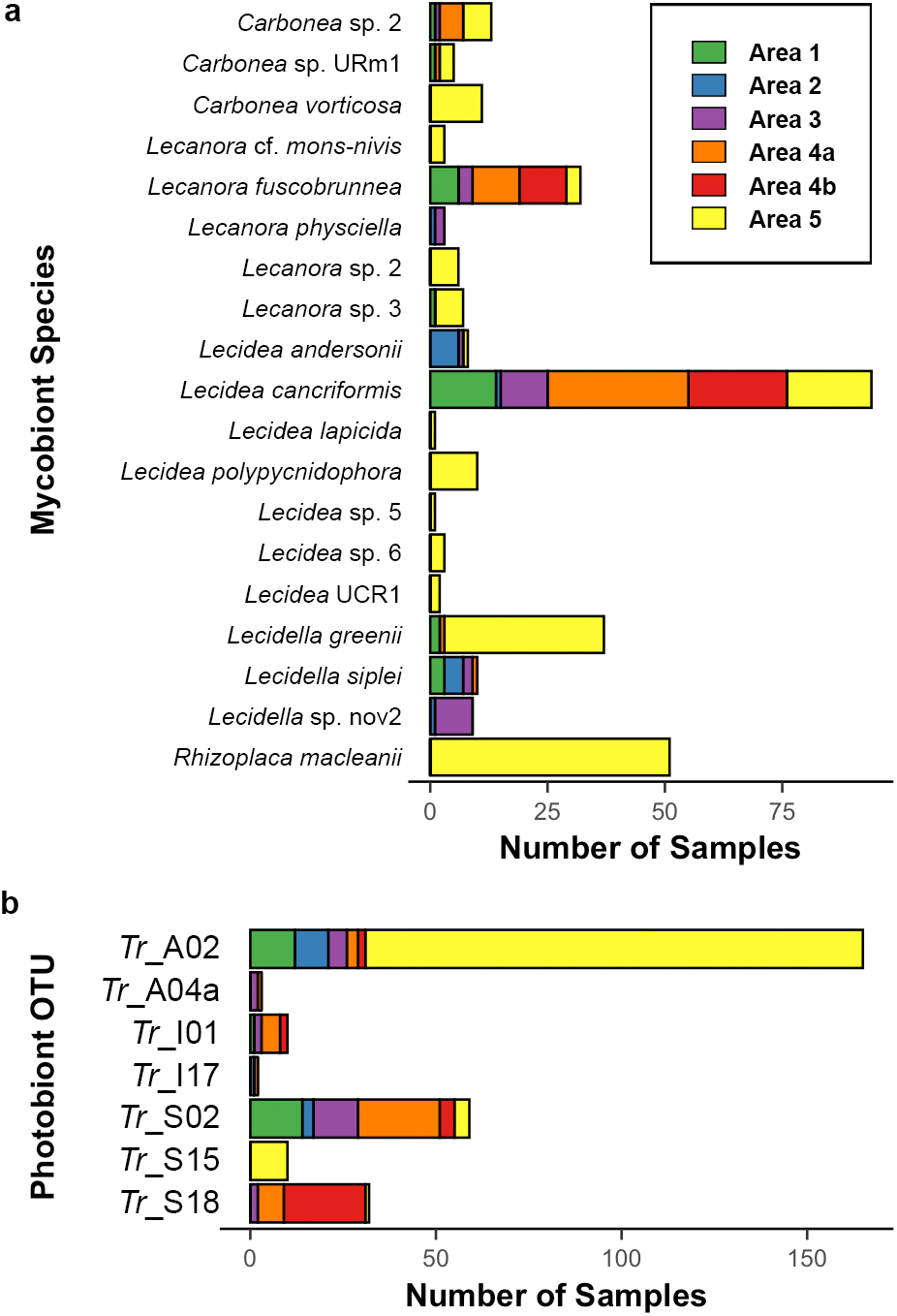
Barplots giving the number of samples per mycobiont species/ photobiont OTU and area included in this study. (a) Mycobiont species (total sample size: n = 306), (b) photobiont OTUs (total sample size: n = 281).

**Supplementary Figure S6.**
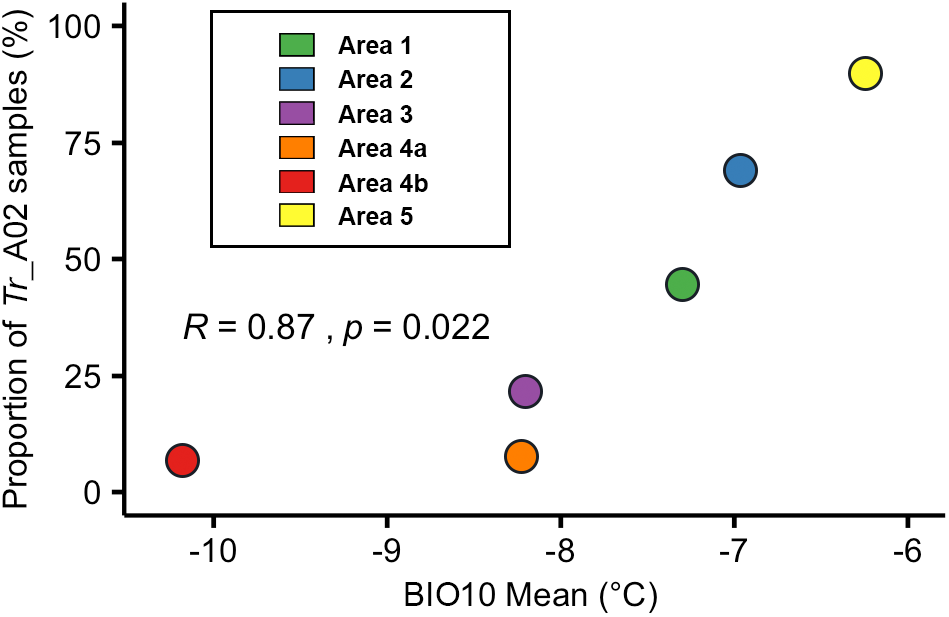
Correlation plot. Percentage of *Trebouxia* OTU A02 samples against mean values of BIO10 (mean temperature of warmest quarter) for the different areas.

**Supplementary Figure S7.**
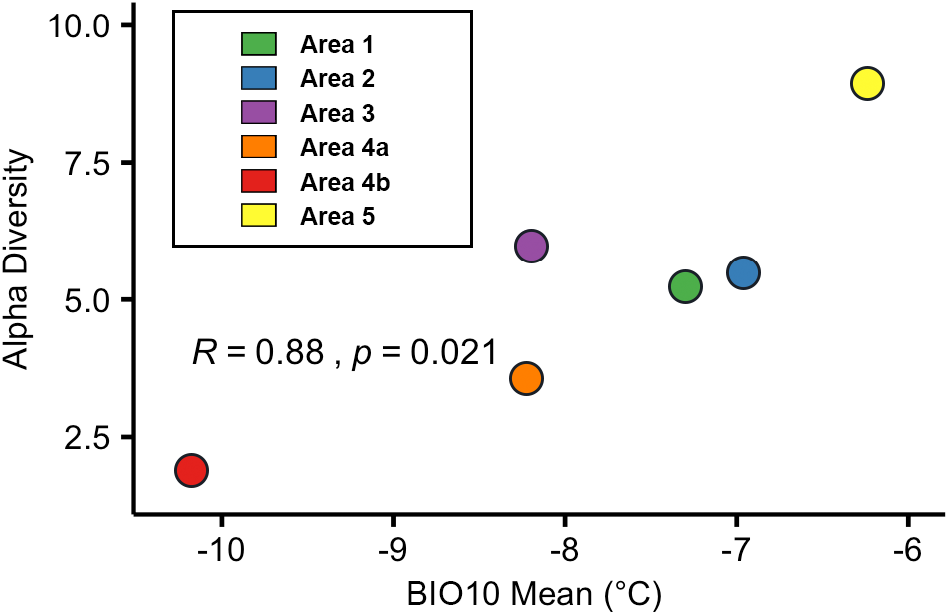
Correlation plot. Alpha diversity values of mycobiont species against BIO10 (mean temperature of warmest quarter) mean values of the different areas.

**Supplementary Figure S8.**
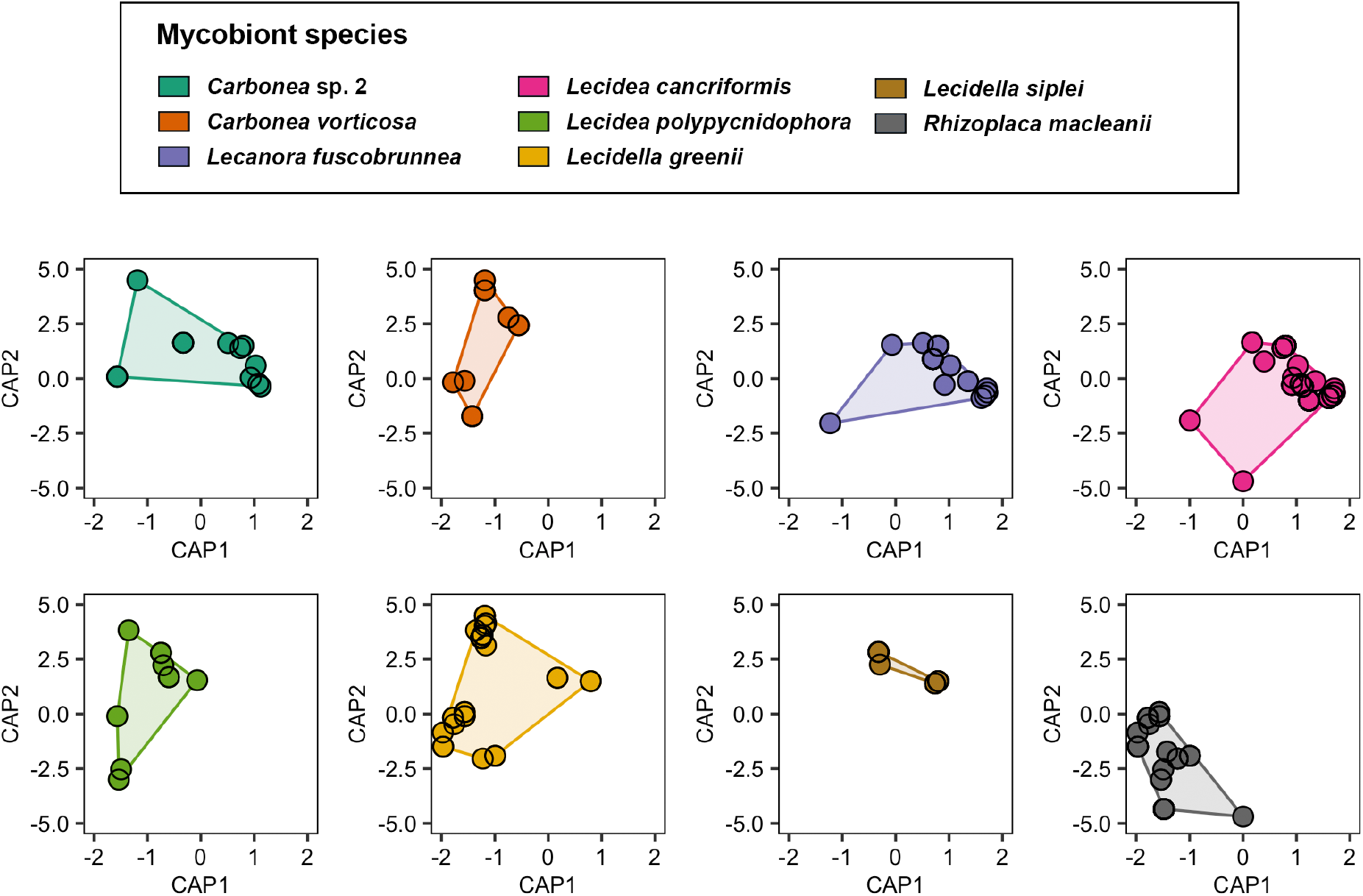
Ordination plots showing the similarity of mycobiont samples with n ≥ 10 after constrained analysis of principal coordinates. Samples located closer to each other are also more similar in terms of the environmental factors elevation, BIO10 and BIO12. The first constrained axis CAP1 explained 13.77 % of the variance, the second constrained axis CAP2 1.76 % of the variance. Only the first axis was significant (F = 17.1640, p = 0.001).

**Supplementary Figure S9.**
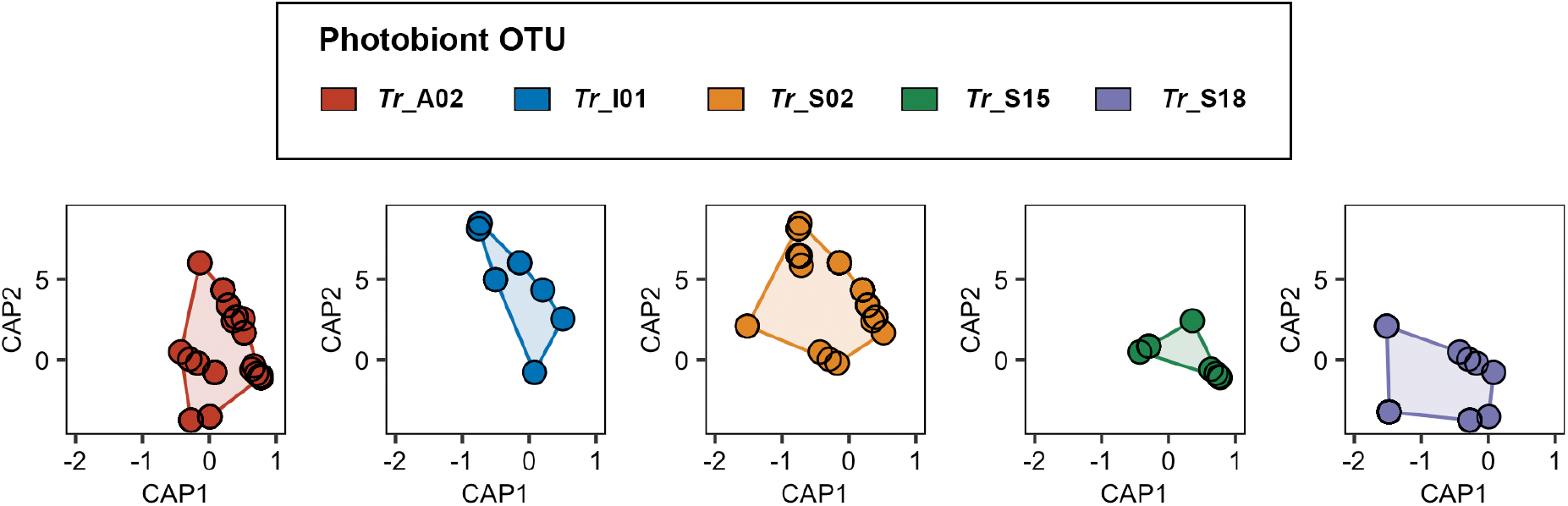
Ordination plots showing the similarity of photobiont OTUS with n ≥ 10 after constrained analysis of principal coordinates. Samples located closer to each other are also more similar in terms of the environmental factors elevation, BIO10 and BIO12. The first constrained axis CAP1 explained 36.87 % of the variance, the second constrained axis CAP2 1.46 % of the variance. Only the first axis was significant (F = 57.0275, p = 0.001).

**Supplementary Figure S10.**
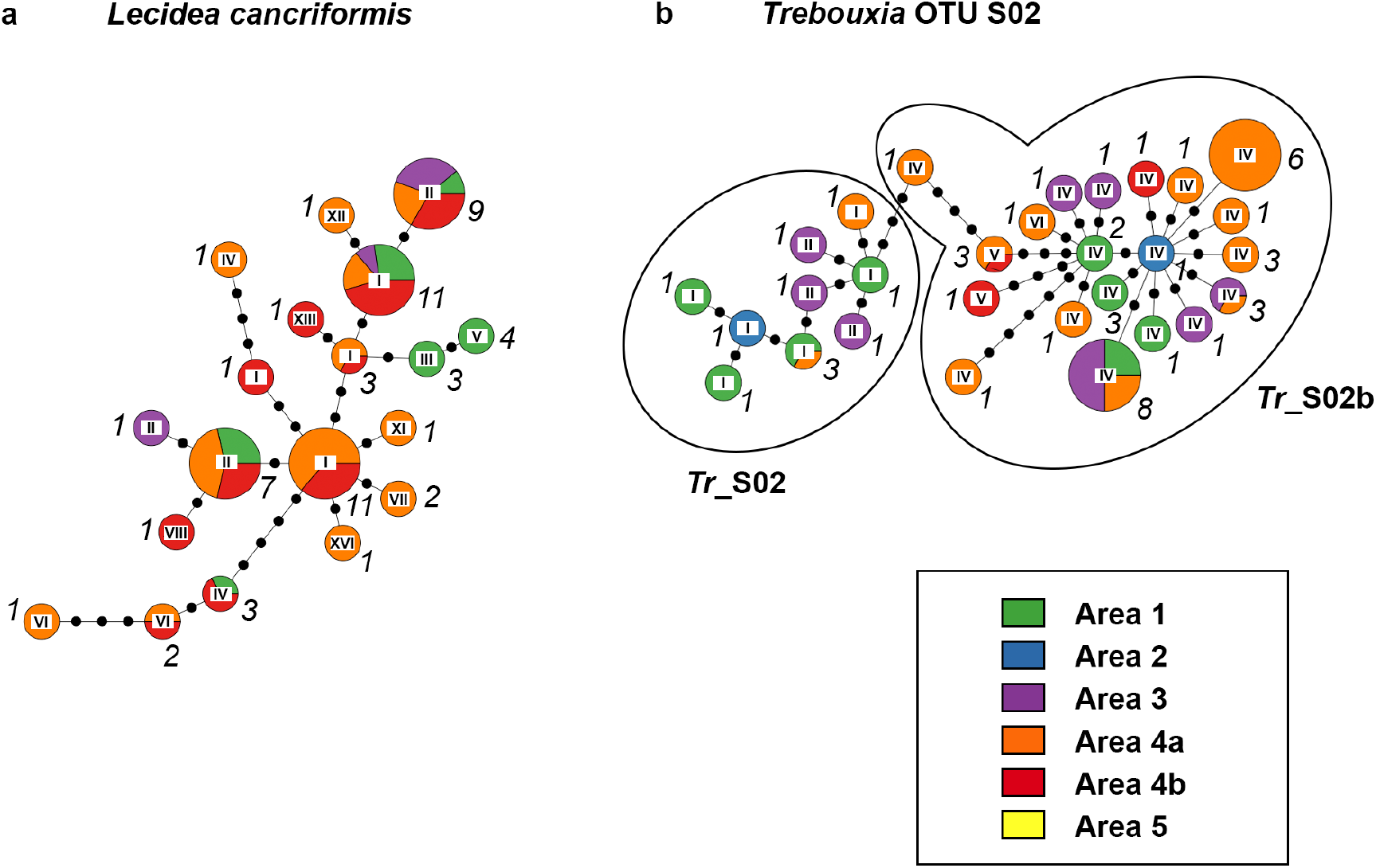
Haplotype networks based on multi-locus sequence data, showing the spatial distribution within the different areas. (a) *Lecidea cancriformis*, (b) *Trebouxia* OTU S02. Roman numerals at the center of the pie charts refer to the haplotype IDs based on ITS data (cf., Fig. 2 and Fig. 3 of main text). The italic numbers next to the pie charts give the total number of samples per haplotype. The circle sizes reflect relative frequency within the species; the frequencies were clustered in ten (e.g. the circles of all haplotypes making up between 20-30 % have the same size).

